# Interaction between host G3BP and viral nucleocapsid protein regulates SARS-CoV-2 replication

**DOI:** 10.1101/2023.06.29.546885

**Authors:** Zemin Yang, Bryan A. Johnson, Victoria A. Meliopoulos, Xiaohui Ju, Peipei Zhang, Michael P. Hughes, Jinjun Wu, Kaitlin P. Koreski, Ti-Cheng Chang, Gang Wu, Jeff Hixon, Jay Duffner, Kathy Wong, Rene Lemieux, Kumari G. Lokugamage, Rojelio E. Alvardo, Patricia A. Crocquet-Valdes, David H. Walker, Kenneth S. Plante, Jessica A. Plante, Scott C. Weaver, Hong Joo Kim, Rachel Meyers, Stacey Schultz-Cherry, Qiang Ding, Vineet D. Menachery, J. Paul Taylor

## Abstract

G3BP1/2 are paralogous proteins that promote stress granule formation in response to cellular stresses, including viral infection. G3BP1/2 are prominent interactors of the nucleocapsid (N) protein of severe acute respiratory syndrome coronavirus 2 (SARS-CoV-2). However, the functional consequences of the G3BP1-N interaction in the context of viral infection remain unclear. Here we used structural and biochemical analyses to define the residues required for G3BP1-N interaction, followed by structure-guided mutagenesis of G3BP1 and N to selectively and reciprocally disrupt their interaction. We found that mutation of F17 within the N protein led to selective loss of interaction with G3BP1 and consequent failure of the N protein to disrupt stress granule assembly. Introduction of SARS-CoV-2 bearing an F17A mutation resulted in a significant decrease in viral replication and pathogenesis in vivo, indicating that the G3BP1-N interaction promotes infection by suppressing the ability of G3BP1 to form stress granules.

## Introduction

Severe acute respiratory syndrome coronavirus 2 (SARS-CoV-2), the virus responsible for the COVID-19 pandemic, uses its viral proteins to manipulate and control host processes. Among the virus’s four structural proteins, the nucleocapsid (N) protein is primarily responsible for compacting viral genomic RNA (gRNA) inside the virion, likely by promoting condensation of gRNA into ribonucleoprotein (RNP) complexes via phase separation ^1,2^. In addition to this primary function, the N protein plays a role in viral RNA replication ^3^, subgenomic RNA transcription ^3–5^, and suppression of innate immune responses ^6^. Given its multiple roles, it is critical to understand how N protein modulates its responses to facilitate SARS-CoV-2 infection. Notably, expression of the SARS-CoV-2 N protein is dependent on the stage of infection: protein levels begin relatively low early in infection and ramp up dramatically as infection proceeds, eventually becoming the most abundant viral protein in infected cells ^7–13^.

Recent work suggests that SARS-CoV-2 N plays a role in the suppression of stress granules, which are large cytoplasmic RNP assemblies that form in response to cellular stresses, including viral infection. The mechanistic connection between N protein and stress granule formation appears to center on the paralogous RNA-binding proteins G3BP1 and G3BP2, which are the most prominent host interactors of the N protein and are key components of the stress granule response ^9,14–19^. G3BP1 is a highly abundant protein in the cytoplasm that surveilles its milieu for a rise in the concentration of uncoated RNAs, such as occurs during stress responses. When such a rise occurs, G3BP1 facilitates condensation of RNAs and proteins to form stress granules ^20^.

While stress granules are generally thought to promote antiviral responses, the relationship between viral infections and G3BP1 is complex. In many types of viral infections, including poliovirus ^21^, coxsackievirus type B3 ^22^, feline calicivirus ^23^, porcine epidemic diarrhoea virus ^24,25^, encephalomyocarditis virus ^26^, foot-and-mouth disease virus ^27^, chikungunya virus ^28^, Semliki Forest virus ^29–32^, enterovirus ^33^, noroviruses ^34^, and Zika virus ^35,36^, viruses neutralize the activity of G3BP1 and inhibit stress granule assembly in order to promote viral replication ^37^. Various strategies have evolved to accomplish this antagonism, including cleavage of the G3BP1 protein ^21–27^ and/or the production of a protein that directly interacts with a key binding pocket in G3BP1 ^28–32^, thereby inhibiting its ability to drive stress granule assembly. Paradoxically, some viruses, such as chikungunya virus ^28^, Semliki Forest ^31^, noroviruses ^34^, Zika Virus ^35,36^, and hepatitis C virus ^38^, also appear to co-opt the condensation properties of G3BP1 to promote their own replication.

Notably, stress granules are nearly absent in SARS-CoV-2-infected cells, despite the presence of double-stranded RNA, which is capable of inducing stress granule formation through PKR activation ^39–42^. Both SARS-CoV-2 infection and ectopic N expression inhibit stress granule formation in response to exogenous stimuli ^17,39–45^. However, the relationship between N protein and G3BP1-dependent stress granule formation in the physiological context of viral infection in vivo are less clear. Indeed, inhibition or loss of G3BP1 has been reported to promote replication in some contexts and cell types ^41,42^ but decrease replication in others ^46,47^.

In this study, we sought to clarify the role of the G3BP1-N interaction in SARS-CoV-2 replication. We began by using structural and biochemical analyses to define the G3BP1-N interaction at the level of individual amino acids, followed by structure-guided mutagenesis of G3BP1 and N to selectively and reciprocally disrupt their interaction. We then used these insights to investigate the impact of disrupting the G3BP1-N interaction in vivo. Our results demonstrate that SARS-CoV-2 directly interacts with host G3BP1 by inserting the side chain of N-F17 into the center of a hydrophobic pocket of the NTF2L domain of G3BP1. This N protein binding competes with other host G3BP1 interactors and blocks the formation of the G3BP1-centered stress granule assembly. Importantly, mutation F17A within the N protein ablates interaction with G3BP1 without disruption of other known G3BP1 interactors. SARS-CoV-2 mutants with the F17A mutation fail to disrupt stress granule assembly, show attenuated viral replication in vitro, and cause reduced disease in vivo. Together, our findings suggest that sequestration of long viral RNAs by stress granules contributes to antiviral defense, and that the G3BP1-N interaction promotes viral infection by suppressing G3BP1-dependent stress granule formation.

## Results

### SARS-CoV-2 N protein directly interacts with G3BP1

Numerous independent analyses have identified proteins that interact with the N protein of SARS-CoV-2 ^9,14–19^. Notably, across diverse cell types, systems, and experimental approaches, G3BP1 and G3BP2 have been consistently observed as the most prominent host interactors of N protein ^9,14–19^. Consistent with these studies, we found G3BP1 and G3BP2 among the top hits in the N protein interactome in our independent mass spectrometry analysis of HEK293T cells expressing GFP-tagged N protein (**Figure 1A, B**). Of all 26 high confidence interactors, six were identified as functioning in protein folding (*HSPA1A*, *HSPA1L*, *HSPA5*, *HSPA6*, *HSPA8*, *HSPA9*), one was identified as being involved in mitochondrial ADP/ATP transport (*SLC25A4*), and the remaining 19 were classified as RNPs involved in various aspects of RNA metabolism, including RNA granule regulation, mRNA splicing, and translation (**Figure 1A**). Remarkably, our analysis of 6 published N protein interactomes, combined with our own dataset, revealed that G3BP1 and G3BP2 were the only proteins consistently identified in all 7 studies (**Figure 1B**).

**Figure 1.**
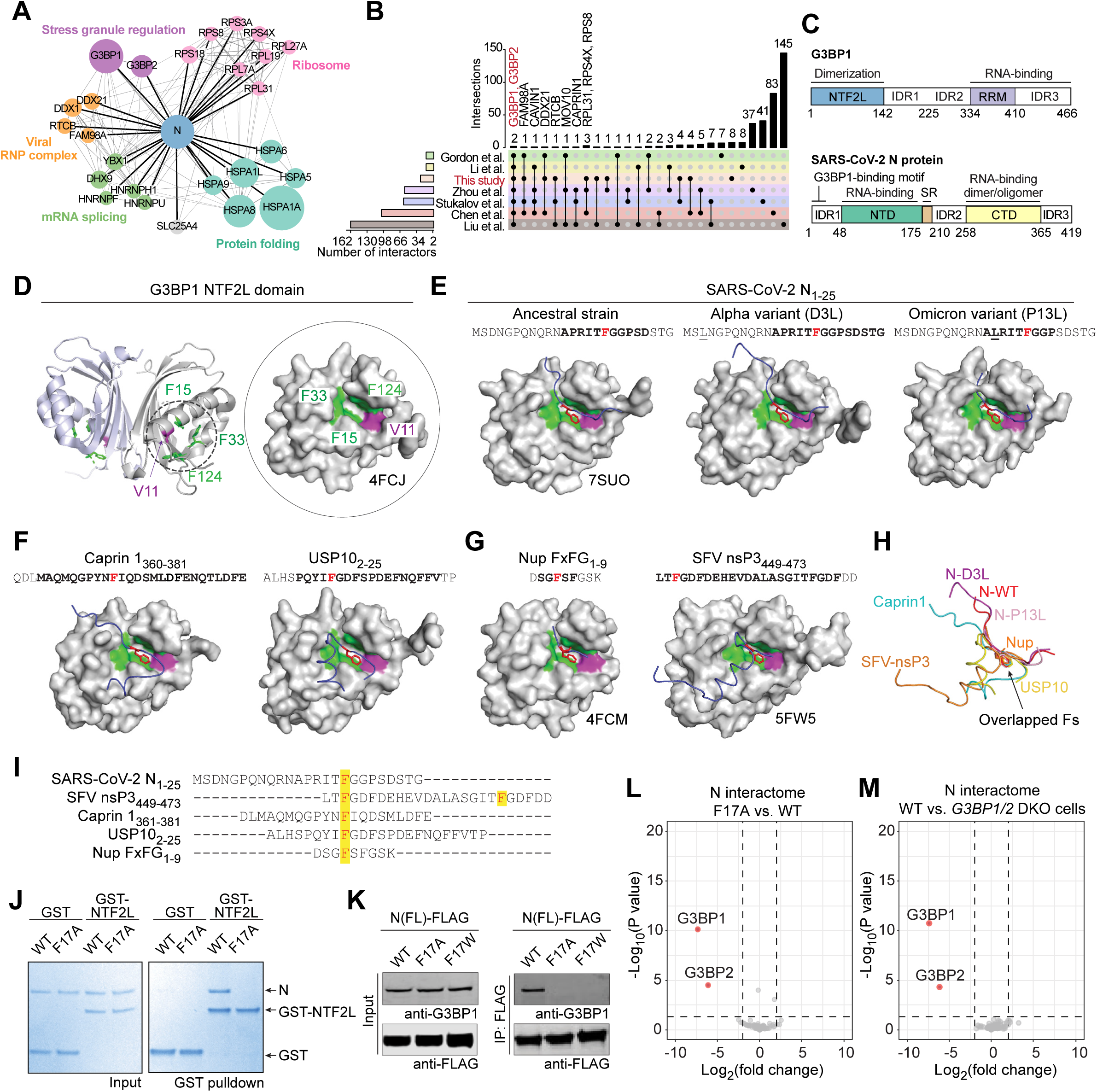
The SARS-CoV-2 N Protein Interacts with Host G3BP1 via N-F17. (A) N protein interactome as defined by mass spectrometry analysis of affinity-purified HEK293T cells expressing GFP-tagged N protein. All 26 high confidence interactors are shown grouped by cellular function. (B) UpSet plot showing comparison of the N interactomes identified in this study and 6 previous studies. G3BP1/2 were the only proteins identified across all 7 studies. (C) Domain organization of G3BP1 and N protein. IDR, intrinsically disordered region. RRM, RNA recognition motif; NTD, N-terminal domain; CTD, C-terminal domain. Residues 1-25 of N protein are highlighted as the G3BP1 binding region. (D) Structure of G3BP1 NTF2L domain (PDB: 4FCJ) with V11, F15, F33, and F124 highlighted in cartoon and surface presentations. Side chains from these 4 residues create a hydrophobic pocket for protein binding. (E) Crystal structures of G3BP1 NTF2L in complex with peptides derived from N-WT_1-25_ (PDB: 7SUO), N-D3L_1-25_, and N-P13L_1-25_. Peptide paths are shown in blue; side chains of N-F17 are shown in red. In the amino acid sequence, F17 is shown in red, mutated residues are underlined, and bold font indicates amino acids traceable in the crystal structures. (F) Crystal structures of G3BP1 NTF2L in complex with peptides derived from caprin 1_361-381_ and USP10_2-25_. Peptide paths are shown in blue; side chains of relevant phenylalanine residues (F372 of caprin 1, F10 of USP10) are shown in red. In the amino acid sequences, F372 and F10 are shown in red and bold font indicates amino acids traceable in the crystal structures. (G) Crystal structures of G3BP1 NTF2L in complex with peptides derived from Nup FxFG_1-9_ (PDB: 4FCM) and SFV nsP3_449-473_ (PDB: 5FW5). Peptide paths are shown in blue; side chains of Nup-FxFG F4 and nsP3 F451 are shown in red. SFV nsP3 F468 also inserted its side chain into the NTF2L binding pocket (not shown). In the amino acid sequences, F4 and F451 are shown in red and bold font indicates amino acids traceable in the crystal structures. (H) Overlay of the paths of all seven NTF2L-interacting peptides as extracted from crystal structures. Side chains are shown for phenylalanine residues that insert into NTF2L. (I) Amino acid sequence of NTF2L-binding peptides; highlights indicate inserted phenylalanines. (J) GST pulldown of purified GST-NTF2L with N-WT and N-F17A. Input (left) and bound proteins (right) are shown. Proteins were visualized with Coomassie Blue staining. (K) HEK293T cells were transfected with FLAG-tagged N-WT, N-F17A, or N-F17W. Cell extracts were captured with magnetic beads conjugated with FLAG antibody for immunoprecipitation and bound proteins were analyzed by immunoblot. (L) The interactomes of N-WT and N-F17A were determined by GFP-tag-based affinity purification from HEK293T cells followed by mass spectrometry. Red and gray dots respectively indicate the interactors that were significantly decreased and unchanged in the N-F17A interactome. The p-value of 0.05 and foldchange of 2 are used as the cutoff. Comparing the two interactomes revealed that G3BP1 and G3BP2 were the only host proteins that showed significant differences. (M) The interactome of N-WT was determined in HEK293T *G3BP1/2* DKO cells as in (L) and compared to the interactome of N-WT in parental HEK293T cells (L). Red and gray dots respectively indicate the interactors that were significantly decreased and unchanged in the *G3BP1/2* DKO cells. The p-value of 0.05 and foldchange of 2 are used as the cutoff. G3BP1/2 were the only significant changes observed for the N interactome between WT and *G3BP1/2* DKO cells.

G3BP1/2 are paralogous proteins that share a common domain architecture consisting of a N-terminal NTF2-like (NTF2L) domain that underlies protein homo-dimerization, two intrinsically disordered regions in the center of the protein (negatively charged and acidic IDR1, slightly positively charged IDR2), and a C-terminal RNA binding domain (RBD) composed of a folded RNA-recognition motif (RRM) in tandem with an arginine-glycine-rich IDR3 (**Figure 1C**). SARS-CoV-2 N protein has two folded domains: an N-terminal RNA-binding domain ^48–50^ and a C-terminal domain that also has RNA binding capacity, exists as a tight dimer in solution, and can further assemble into oligomers in vitro ^50–54^. These two folded domains are flanked by three intrinsically disordered regions: IDR1 (also known as N-arm), IDR2, which is a central serine-arginine-rich flexible linker region (also known as LKR), and IDR3 (also known as C-tail) (Figure 1C) ^55,56^.

We next examined whether the G3BP1-N protein interaction was direct, and if so, which domains mediated this interaction. To this end, we used in vitro pulldown assays with purified N (His-SUMO-tagged) and G3BP1 (GST-tagged) proteins. We found that His-SUMO-N and GST-G3BP1 could each pull down the other protein, indicating a direct interaction (**Supplementary Figure 1A**). To test whether this interaction was mediated by the G3BP1 NTF2L domain, we repeated these experiments using purified GST-NTF2L as well as full-length GST-G3BP1 ΔNTF2L. The NTF2L domain alone was able to bind the N protein, whereas the protein lacking the NTF2L domain failed to do so, indicating that the NTF2L domain of G3BP1 is necessary and sufficient for interaction with the N protein (**Supplementary Figure 1A**).

To identify the G3BP1 binding site within the N protein, we generated a series of N protein truncations with sequential domain deletions (**Supplementary Figure 1B**), which were then transfected into HEK293T cells and tested for their ability to immunoprecipitate endogenous G3BP1 (**Supplementary Figure 1B-C**). We found that deletion of N-IDR1 (aa 1-47) was sufficient to abolish the interaction of N protein with G3BP1, whereas deletion of IDR2, IDR3, or the C-terminal domain had relatively little effect (**Supplementary Figure 1B, C**). We then further mapped the G3BP1 binding site using serial deletions within IDR1, finding that the first 25 amino acids were sufficient for a robust interaction with G3BP1 (**Supplementary Figure 1D**). Taken together, these results demonstrate a direct interaction between the N protein and G3BP1 that requires the NTF2L domain of G3BP1 and the first 25 amino acids of the N protein, consistent with previous studies ^43,46,57^.

### N protein and G3BP1 interactors bind to G3BP1 NTF2L in a mutually exclusive manner

Whereas the G3BP1-N protein interaction has been shown to occur within the NTF2L domain, the effect of this binding on the normal G3BP1 host interaction network has not yet been explored. The NTF2L domain of G3BP1 harbors the binding site for several types of proteins, including host proteins that regulate stress granule assembly (e.g., caprin 1 ^58^ and USP10 ^59^) as well as viral proteins (e.g., non-structural protein 3 (nsP3) of alphaviruses ^60–62^). Several of these binding partners are known to bind G3BP1 in a mutually exclusive manner. For instance, caprin 1 and USP10 show mutually exclusive binding to the NTF2L domain, with antagonistic effects on stress granule formation ^63^. To determine whether these interactors compete with N protein for binding to NTF2L, we synthesized three short peptides derived from the G3BP1 interacting motifs from caprin 1 (aa 361-381), USP10 (aa 1-25), and nsP3 (aa 449-473) of the alphavirus Semliki Forest virus (SFV) (**Supplementary Figure 1E**). We then used an in vitro GST pulldown assay with purified GST-NTF2L and His-SUMO-N proteins and added increasing concentrations of caprin 1, USP10, or nsP3 peptides to test the effect of these peptides on the NTF2L-N interaction. As expected, all three of these peptides competed against the NTF2L-N interaction in a dose-dependent manner, with the nsP3-25mer peptide showing the most effective competition for binding to N protein (**Supplementary Figure 1F**). We then tested for competitive binding in living cells by expressing nsP3-25mer-mCherry and G3BP1-GFP in *G3BP1/2* double knockout (DKO) cells. We found that the interaction between GFP-G3BP1 and N protein was dramatically reduced in the presence of nsP3-25mer (**Supplementary Figure 1G**), demonstrating that this mutually exclusive binding can also occur in the context of living cells.

### A common docking mode: N and other G3BP1 interactors insert a phenylalanine side chain into the center of a NTF2L hydrophobic pocket

The mutually exclusive binding by N protein and other G3BP1 interactors suggested that these proteins may bind to the same site on NTF2L. However, the primary amino acid sequences of G3BP1-interacting motifs look quite different from each other and lack a conserved motif across all NTF2L interactors (**Figure 1I**). To examine this question more deeply, we turned to crystallography to examine whether a shared interaction mode might exist at the 3-dimensional level.

Published structures of G3BP1/2 have revealed that the NTF2L domain consists of a five-stranded antiparallel beta-sheet and three alpha-helices (**Figure 1D**) ^64^. Two NTF2L molecules form a homodimer by the face-to-face stacking of two beta-sheets. On the outer surface of each dimer, two alpha-helices, with side chains of residues V11, F15, F33, and F124 (**Figure 1D**, green and purple surfaces), create a hydrophobic pocket for protein binding ^64^. This pocket is responsible for the vast majority of G3BP1 protein interactions relative to the formation of a condensation network (unpublished) ^20^.

The crystal structure of the G3BP1 NTF2L domain in complex with the first 25 amino acids of the N protein (N-WT_1-25_) has been reported ^65^. In this structure, the side chain of N-F17 (**Figure 1E**, red highlight) was inserted into the center of the NTF2L pocket and also interacted with surrounding NTF2L residues. To compare this NTF2L-N interaction with other NTF2L interactors, we obtained the crystal structures of G3BP1 NTF2L in complex with peptides derived from two host proteins, caprin 1_360-381_ and USP10_2-25_ (**Figure 1F**). Similar to SARS-CoV-2 N protein and consistent with published structures ^58,59^, both peptides inserted one phenylalanine (F372 in caprin 1, F10 in USP10) into the same NTF2L pocket. This finding was consistent with published structures of NTF2L interacting with an FxFG peptide derived from host nucleoporin (Nup) protein (PDB: 4fcm) ^64^ and nsP3 protein of SFV (PDB: 5FW5) ^61^, where phenylalanines (F4 in Nup FxFG peptide, F451 and F468 in nsP3) were inserted into this pocket (**Figure 1G**).

Notably, two SARS-CoV-2 variants of concern ^66–68^, Alpha and Omicron, have acquired mutations within the first 25 amino acids of the N protein, though neither has been examined structurally. To test whether these mutations might alter the G3BP1-N protein interaction, we determined the crystal structures of G3BP1 NTF2L in complex with N-D3L_1-25_ (Alpha variant mutation ^69^) and N-P13L_1-25_ (Omicron variant mutation ^70^) (**Figure 1E**). Comparing these two structures with NTF2L bound to N-WT_1-25_, we found that all three N peptides bound to the same pocket of NTF2L by inserting the side chain of F17 (**Figure 1E**, red highlight) into the center of this pocket. N-D3L could not be visualized in the structure, indicating that this residue is located some distance from the NTF2L-N interaction surface (**Figure 1E**). N-P13L, although in close proximity to the interacting interface, was determined to be not essential for the interaction (**Figure 1E**). Interestingly, N-P13L slightly altered the orientation of the peptide in the proximity of the NTF2L interacting site, raising the possibility of alterations in binding between NTF2L and N-P13L, a prediction we pursue in greater detail below. Nonetheless, we found that the G3BP1-N interaction was essentially unaltered even after the emergence of SARS-CoV-2 mutants. Indeed, after extracting and overlaying the structures of all NTF2L-interacting peptides, we found that the side chain of the inserted phenylalanines of different peptides adopt a nearly identical conformation (**Figure 1H, I**).

We next tested these structural observations in cells by mutating relevant phenylalanine residues in peptides derived from NTF2L interactors and assessing the ability of these peptides to bind endogenous G3BP1. Consistent with our structural predictions, substituting phenylalanines with alanines in caprin 1 (F372A), USP10 (F10A), nsP3 (F451A/F468A), or SARS-CoV-2 N (F17A) abolished their interaction with endogenous G3BP1 in HEK293T cells (**Supplementary Figure 1H**). We also tested the effect of F17A and F17W substitutions in full-length N protein, finding that both F17A and F17W abolished the interaction of N protein with G3BP1 both in vitro (**Figure 1J**) and in cells (**Figure 1K**).

### N-F17A specifically abolishes interaction with G3BP1/2

To examine the consequences of the SARS-CoV-2 N-F17A mutation on the interaction of N protein with other host proteins, we compared the interactome of N-WT and N-F17A in HEK293T cells by GFP-tag-based affinity purification and mass spectrometry. Remarkably, the only host proteins that showed significant changes across duplicate experiments were G3BP1 and G3BP2 (**Figure 1L**), suggesting that N-F17 is specifically utilized for interaction with G3BP1/2 in host cells. We next considered whether G3BP1/2 might also serve as an adaptor for the N protein to recruit secondary interactors. To test this idea, we also defined the interactome of the N protein in HEK293T *G3BP1/2* DKO cells. G3BP1/2 were once again the only significant changes observed for the N interactome between wild-type (WT) and *G3BP1/2* DKO cells, consistent with our hypothesis that G3BP1/2 do not act as intermediates to recruit other interactors to the N protein (**Figure 1M**).

### N-F17A does not alter the in vitro phase separation behavior of N protein

During viral infections, the N protein engages in both homotypic interactions with other N proteins and heterotypic interactions with RNA molecules. These interactions underlie the ability of the N protein to undergo liquid-liquid phase separation ^1^. To test whether F17 affects these multivalent interactions, we assessed the ability of purified recombinant N-WT and N-F17A to undergo phase separation in vitro. We found that phase separation of N protein triggered by the addition of crowding agent (**Supplementary Figure 2A**) or RNA (**Supplementary Figure 2B**) was unaltered between N-WT and N-F17A, demonstrating that N-F17A does not disrupt heterotypic interactions with RNA or homotypic interactions with other N proteins. We further assessed the ability of N-WT and N-F17A to bind RNA in cells using cross-linking and immunoprecipitation (CLIP), finding that N-WT and N-F17A bind similarly to cellular RNAs (**Supplementary Figure 2C**).

### G3BP1-N protein interaction is conserved among all SARS-CoV-2 variants

To determine whether the G3BP1-N protein interaction is conserved among SARS-CoV-2 variants, we analyzed the amino acid variations of N protein among 8,297,154 viral genomes deposited in GISAID by March 2022 ^71^. Most amino acids within the protein were highly conserved (**Figure 2A**). F17 showed no amino acid variations among all viral genomes (**Figure 2A**) and was conserved in all viral lineages, including transient sublineages that have appeared and declined over the course of the pandemic (**Figure 2B**). For the other amino acids within aa 1-25, the region that is both necessary and sufficient for G3BP1 binding, most amino acids were also highly conserved. Exceptions here were three amino acid replacements, D3L, Q9L, and P13L – two of which we examined above using crystallography – which were respectively observed in 13.3%, 3.4%, and 22.3% of all SARS-CoV-2 genomes (**Figure 2A**). D3L appeared specifically in the Alpha variant, whereas Q9L was observed in a sublineage of the Delta variant. Interestingly, P13L appeared sporadically among early variants in 2020 (lineages 20A, 20B, 20C, 20D by Nextstrain ^72^), disappeared in Alpha, Gamma, and Delta variants, and then re-appeared in the Omicron variant and its sublineages (**Figure 2C**).

**Figure 2.**
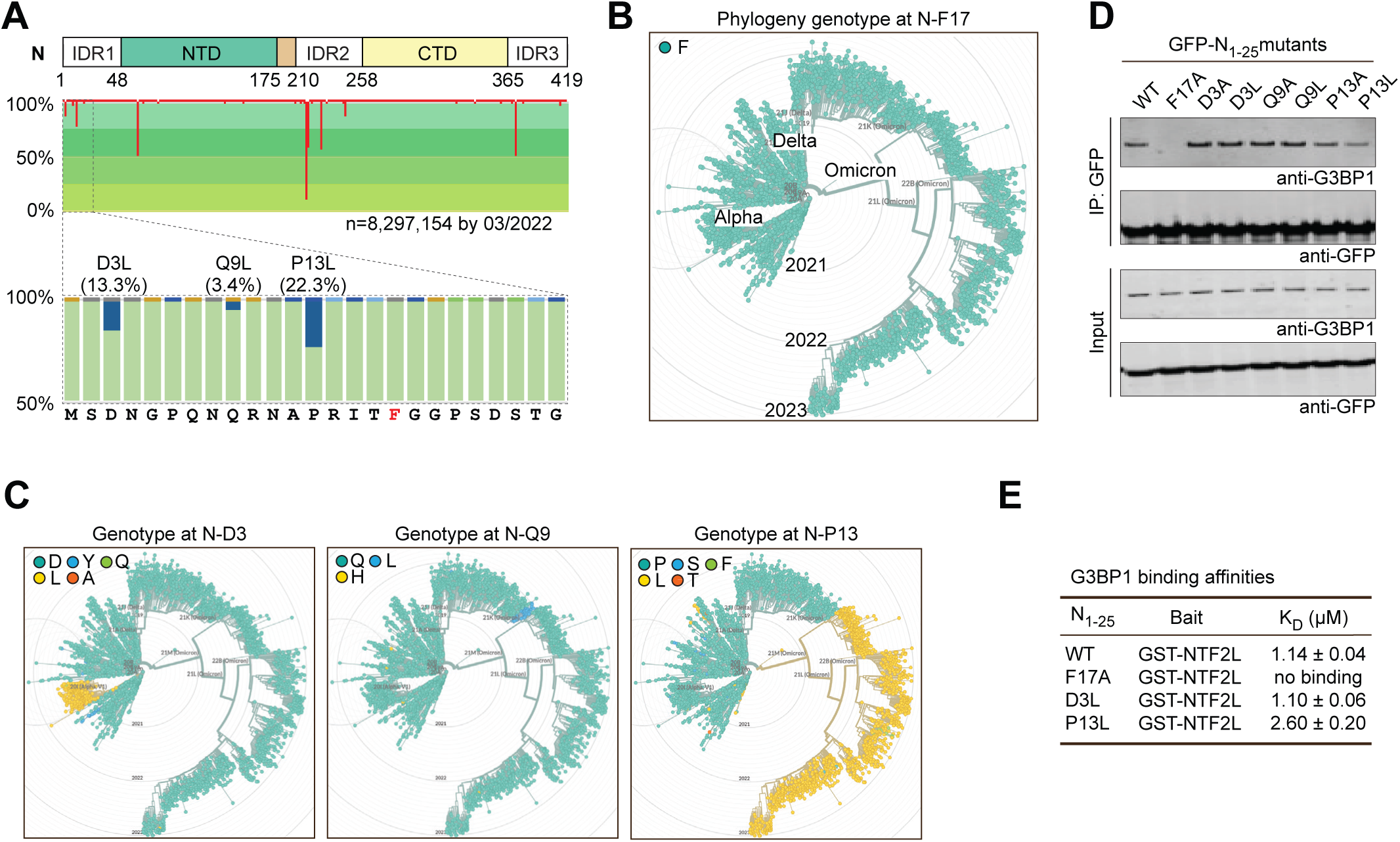
The G3BP1-N Interaction is Conserved Among SARS-CoV-2 Variants. (A) Amino acid variations of N protein among 8,297,154 SARS-CoV-2 genomes deposited in GISIAD by March 2022. Sites with amino acid replacements are highlighted by red lines, with line length indicating the percentage of amino acid variations at that particular site. The conservation of each amino acid is represented by different shades of green. D3L, Q9L, and P13L were observed in 13.3%, 3.4%, and 22.3% of all SARS-CoV-2 genomes, respectively. F17 (red font) showed no amino acid variations. (B, C) Phylogeny genotypes of N protein at F17 (B), D3, Q9, and P13 (C) are shown in chronological order from 2019 to 2023. F17 was conserved in all viral lineages, whereas D3L appeared specifically in the Alpha variant, Q9L appeared in a sublineage of the Delta variant, and P13L appeared sporadically among early variants, disappeared in Alpha, Gamma, and Delta variants, and re-appeared in the Omicron variant and its sublineages. (D) HEK293T cells were transfected with GFP-tagged N_1-25_ with indicated mutations. Cell extracts were captured with magnetic beads conjugated with GFP antibody for immunoprecipitation and bound proteins were analyzed by immunoblot. (E) Results from SPR assay showing binding affinities of indicated N protein peptides with purified GST-NTF2L in vitro.

To determine whether the interaction between N protein and G3BP1/2 was affected by these mutations, we expressed N protein with mutations at each of these residues (D3A, D3L; Q9A, Q9L; P13A, P13L) in HEK293T cells and tested their interaction with endogenous G3BP1 by co-IP. All tested mutants retained the ability to bind G3BP1 (**Figure 2D**), consistent with our earlier crystal structure data in which N-D3L and N-P13L peptides formed a stable complex with NTF2L (**Figure 1E**). Interestingly, N-P13L and N-P13A showed slightly decreased interaction with G3BP1 (**Figure 2D**). We confirmed this observation using surface plasmon resonance (SPR), which revealed a lower binding affinity of N_1-25_-P13L peptide (K_D_ = 2.6 μM) with purified GST-NTF2L protein in vitro, compared to N_1-25_-WT (K_D_ = 1.14 μM) and N_1-25_-D3L (K_D_ = 1.10 μM) peptides (**Figure 2E**). Taken together, these data demonstrate the conservation of the G3BP1-N interaction across all SARS-CoV-2 variants, suggesting that this interaction may be of significance for the virus.

### G3BP1 and N protein interaction specifically involves V11 and F124 of G3BP1

Although all of the NTF2L-interacting peptides that we examined rely on a key phenylalanine to interact with G3BP1 (**Figure 1** and **Supplemental Figure 1**), the amino acids surrounding this phenylalanine residue differ across the various NTF2L interactors, suggesting that the docking modes of these interactors on NTF2L are not identical. In caprin 1, USP10, and SFV-nsP3, we observed that amino acids around the key phenylalanine residue were in close proximity to a side of the NTF2L pocket that is made up of G3BP1 F33 and surrounding amino acids (**Figure 3A**). In contrast, the amino acids surrounding F17 on the N protein were in close contact with the opposing side of the NTF2L binding pocket, which has as its central components G3BP1 F124 and G3BP1 aa 1-11 (**Figure 3B**). These structural insights led us to hypothesize that it would be possible to engineer specific disruption of the G3BP1-N interaction without altering the interaction of G3BP1 with other host proteins.

**Figure 3.**
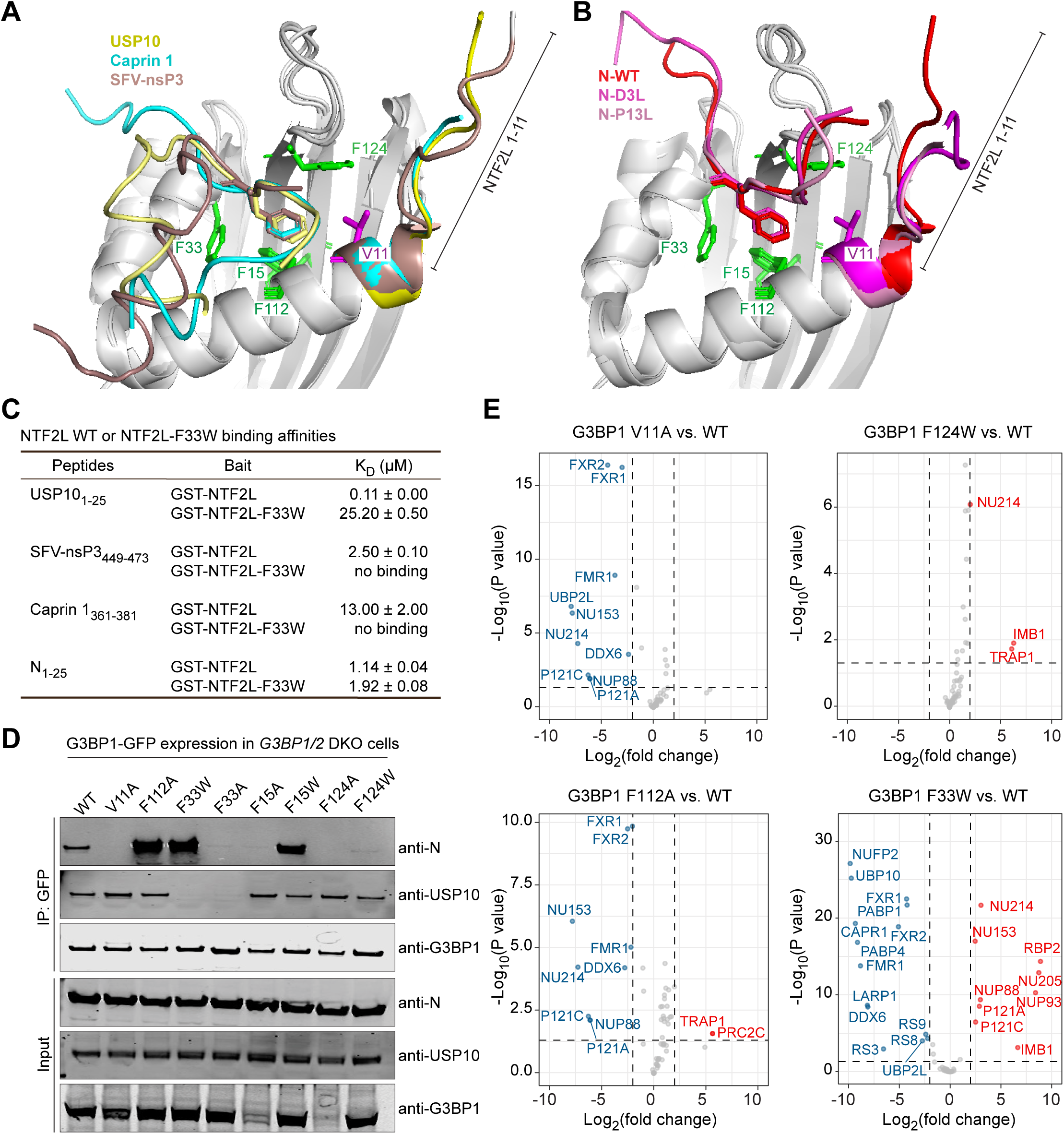
The G3BP1-N Interaction Involves V11 and F124 of G3BP1 NTF2L. (A-B) Alignment of crystal structures of G3BP1 NTF2L bound to USP10_2-25_, caprin 1_361-381_, or nsP3-SFV_449-473_ (PDB: 5FW5) (A) or bound to N-WT_1-25_ (7SUO), N-D3L_1-25_, and N-P13L_1-25_ (B). The N-terminal tail (aa 1-11) of G3BP1 is labeled; side chains of G3BP1 F15, F33, F112, F124, and V11 are labeled and colored green (phenylalanines) or magenta (valine). (C) Results from SPR assay showing binding affinities of indicated peptides with purified GST-NTF2L-WT and GST-NTF2L-F33W proteins. (D) HEK293T *G3BP1/2* DKO cells were co-transfected with N-FLAG and indicated G3BP1-GFP constructs. Cell extracts were captured with magnetic beads conjugated with GFP antibody for IP and bound proteins were analyzed by immunoblot. (E) Comparison of the interactomes of G3BP1-WT with indicated G3BP1 mutants in HEK293T *G3BP1/2* DKO cells. Red, blue, and gray dots respectively indicate the interactors that were significantly increased, decreased, and unchanged in the *G3BP1/2* DKO cells. The p-value of 0.05 and foldchange of 2 are used as the cutoff.

We began by examining the effect of mutating G3BP1 F33, the central amino acid within the NTF2L binding pocket used by caprin 1, USP10, and SFV-nsP3. By SPR, we found that NTF2L-F33W abolished the binding of caprin 1_361-381_, USP10_1-25_, and nsP3_449-473_ peptides to NTF2L, but had relatively little effect on the interaction between N_1-25_ and NTF2L, as evidenced by only a minor decrease in binding affinity (K_D_ = 1.14 μM for NTF2L-WT; K_D_ = 1.92 μM for NTF2L-F33W) (**Figure 3C**). Pursuing this result, we introduced a series of single point mutations to impair the different surfaces of the NTF2L hydrophobic pocket, including V11, F15, F33 and F124, and F112, the phenylalanine very close to the binding surface. When we expressed these G3BP1-GFP mutants along with FLAG-tagged N protein in *G3BP1/2* DKO cells and assessed their interaction by co-IP, we found that G3BP1 V11A and F124A abolished the G3BP1-N interaction, and G3BP1 F124W, F15A, and F33A dramatically reduced the G3BP1-N interaction (**Figure 3D**). In contrast, G3BP1 F112A, F15W, and F33W showed increased G3BP1-N interaction (**Figure 3D**). These findings are consistent with a previous study in which G3BP1 NTF2L domain with V11A, F15A, or F124A mutations lost the interaction with N peptide in vitro ^65^.

Taking stock of these findings and seeking to identify point mutations specifically targeting the N protein interaction, we excluded G3BP1 F33A/F33W because these mutations also caused the loss of interaction of USP10 (**Figure 3D**), as well as G3BP1 F15A and F124A because these mutant proteins expressed at much lower levels, reflecting potential issues with protein stability. We selected the three remaining mutations (V11A, F112A, F124W) for further unbiased analysis by mass spectrometry. For G3BP1 V11A, we found that most of the 58 high-confidence interactors remained intact compared with G3BP1 WT. Notable exceptions included the loss of UBP2L and four Nups that harbor an FxFG motif (Nup153, Nup214, Nup88, Nup121), as well as a reduced interaction with FMR1 and FXR1/2 (**Figure 3E**). To our surprise, G3BP1 F112A lost the interaction with these same four FxFG-containing Nups, but not the other proteins affected by the V11A mutation. The interactome of G3BP1 F124W was overall similar to G3BP1 WT (**Figure 3E**).

Taken together, these studies identify three G3BP1 point mutations that alter the G3BP1-N interaction, albeit with different degrees of efficacy and specificity. G3BP1 V11A abolished the interaction with N protein and impaired the interaction with 7 of 58 G3BP1 interactors. G3BP1 F124W was not as potent as V11A in impairing the interaction with N protein, but retained largely the same interactome as G3BP1 WT. In contrast, G3BP1 F112A increased the interaction with N protein, but also lost interaction with 4 nucleoporin proteins.

### N-F17A reduces SARS-CoV-2 replication in a cell type-dependent manner

After charactering the interaction between N protein and G3BP1 in detail, we introduced these mutations to a virus-like particle or authentic SARS-CoV-2 virus to evaluate the consequences of loss of G3BP1-N interaction on SARS-CoV-2 replication. We first tested the effect of N-F17A, the mutation that specifically abolishes interaction with G3BP1/2, on infection. To this end, we employed our N protein trans-complementation system (**Figure 4A**) ^73^ in which the *N* gene of the SARS-CoV-2 genome is replaced by a GFP reporter (SARS-CoV-2 GFP/ΔN) and the N protein is supplied in trans in Caco-2 cells (Caco2-N cells) to produce replication-competent virus-like-particles (trVLPs) ^43^. In this study, we infected both Caco2-N (WT) and N (F17A) cells with trVLPs at an MOI of 0.05 for 36 hours. The infectivity of newly reproduced trVLPs from Caco2-N (F17A) cells was significantly reduced compared to Caco2-N (WT) cells, as indicated by a >50% decrease in the percentage of GFP-positive cells among cells inoculated by the cell culture supernatant (**Figure 4B**). These results indicate that mutation N-F17A attenuates viral infection in this system.

**Figure 4.**
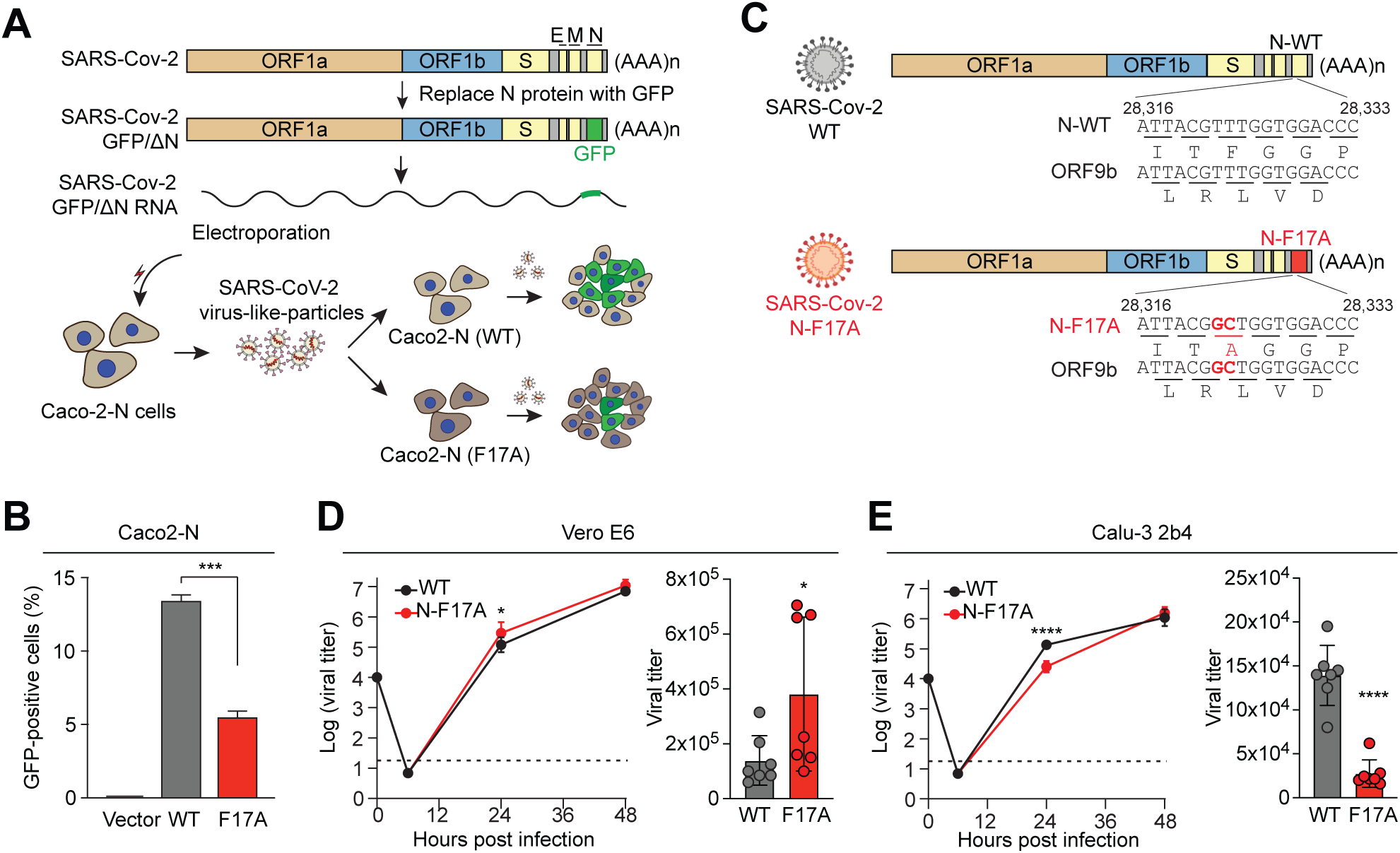
N-F17A Reduces SARS-CoV-2 Replication in Caco2 and Calu-3-2b4 Cells but Not in Vero E6 Cells. (A) Illustration of the strategy for comparing the reproduction of SARS-CoV-2 GFP/ΔN trVLP in Caco2-N (WT) and Caco2-N (F17A) cells. SARS-CoV-2 GFP/ΔN RNA was transcribed in vitro and introduced into Caco2-N cells to produce infectious SARS-CoV-2 trVLPs. Both Caco2-N (WT) and Caco2-N (F17A) cells were infected by SARS-CoV-2 trVLPs at MOI of 0.05 for 36 h. The newly reproduced trVLPs in the supernatant of both cell lines were collected to infect fresh Caco2-N cells for another 36 h. (B) Infected cells (as described in A) were analyzed by flow cytometry to measure the percentage of GFP-positive cells. Error bars represent mean ± SD. ***P < 0.001 by one-way ANOVA with post-hoc Turkey test. (C) Illustration showing introduction of N-F17A into SARS-CoV-2 genome by reverse genetics. Open reading frames of both N and ORF9b are shown. (D-E) Vero E6 (D) and Calu-3 2b4 (E) cells were infected with WT or mutant SARS-CoV-2 (N-F17A) at MOI of 0.01. Viral titers were determined at 0, 6, 24 and 48 hpi. Error bars represent mean ± SD. \**P* < 0.05; \*\*\*\**P* < 0.0001 by two-tailed Student’s t test.

To further pursue this result, we next used our SARS-CoV-2 reverse genetic system ^74,75^ and introduced the N-F17A mutation into the Washington-1 (WA-1) strain (**Figure 4C**). Introduction of N-F17A mutation resulted in a replication-competent virus with no deficit in stock generation or change in foci morphology. After recovery of recombinant SARS-CoV-2, we measured the replication kinetics of the N-F17A mutant relative to WT SARS-CoV-2 in Vero E6 cells ^76,77^, an immortalized cell line established from kidney epithelial cells of an African green monkey, and in Calu-3 2b4 cells ^78^, a human respiratory cell line. Briefly, cells were infected with WT or the N-F17A mutant at an MOI of 0.01 and viral titer was determined over a 48-hour time course. In Vero E6 cells, the N-F17A mutant slightly enhanced replication relative to WT virus at 24 hours post infection (hpi) (**Figure 4D**). In contrast, the N-F17A mutant significantly reduced viral replication by >8-fold at 24 hpi, indicating that the N-F17A mutation attenuates SARS-CoV-2 in relevant human respiratory cells in vitro (**Figure 4E**). Similarly inconsistent results have also been obtained in a variety of in vitro systems designed to test the role of G3BP1 in viral replication ^41,42,46,47^, likely reflecting the challenges associated with using simple in vitro models that lack the complexity of physiological systems.

### N-F17A reduces SARS-CoV-2 replication and pathogenesis in vivo

To help resolve these inconsistent in vitro results, we moved to an in vivo model infection using Golden Syrian hamsters. As previously described ^79^, three- to four-week-old male Golden Syrian hamsters were intranasally infected with either PBS alone (mock), WT, or N-F17A SARS-CoV-2 and monitored for disease for 7 days post infection (dpi) (**Figure 5A**). At 2 and 4 dpi, cohorts of animals underwent nasal washing followed by euthanasia and harvesting of lung tissue to assay viral replication and lung pathology. Notably, N-F17A-infected animals had reduced weight loss relative to WT-infected animals throughout the experimental time course, demonstrating reduced pathogenesis (**Figure 5B**). Though the N-F17A mutant had no effect on viral titer in the nasal washes, N-F17A-infected animals had significantly lower viral titer in the lungs at 2 dpi compared with WT-infected animals (**Figures 5C, D**). In addition, histopathological analysis of lung tissue revealed a significant reduction in overall pathology in N-F17A-infected animals compared with WT-infected animals at 4 dpi (**Figure 5E**). Compared with mock infection, WT-infected animals exhibited widespread interstitial pneumonia. Mononuclear cell infiltration of alveolar septa was observed with numerous enlarged cytopathic alveolar pneumocytes including multinucleation and prominent nucleoli (**Figure 5F**). In contrast, N-F17A-infected animals exhibited patchy interstitial pneumonia with few cytopathic alveolar pneumocytes (**Figure 5F**). Together, these data demonstrate that the N-F17A mutation attenuates SARS-CoV-2 replication and pathogenesis in vivo.

**Figure 5.**
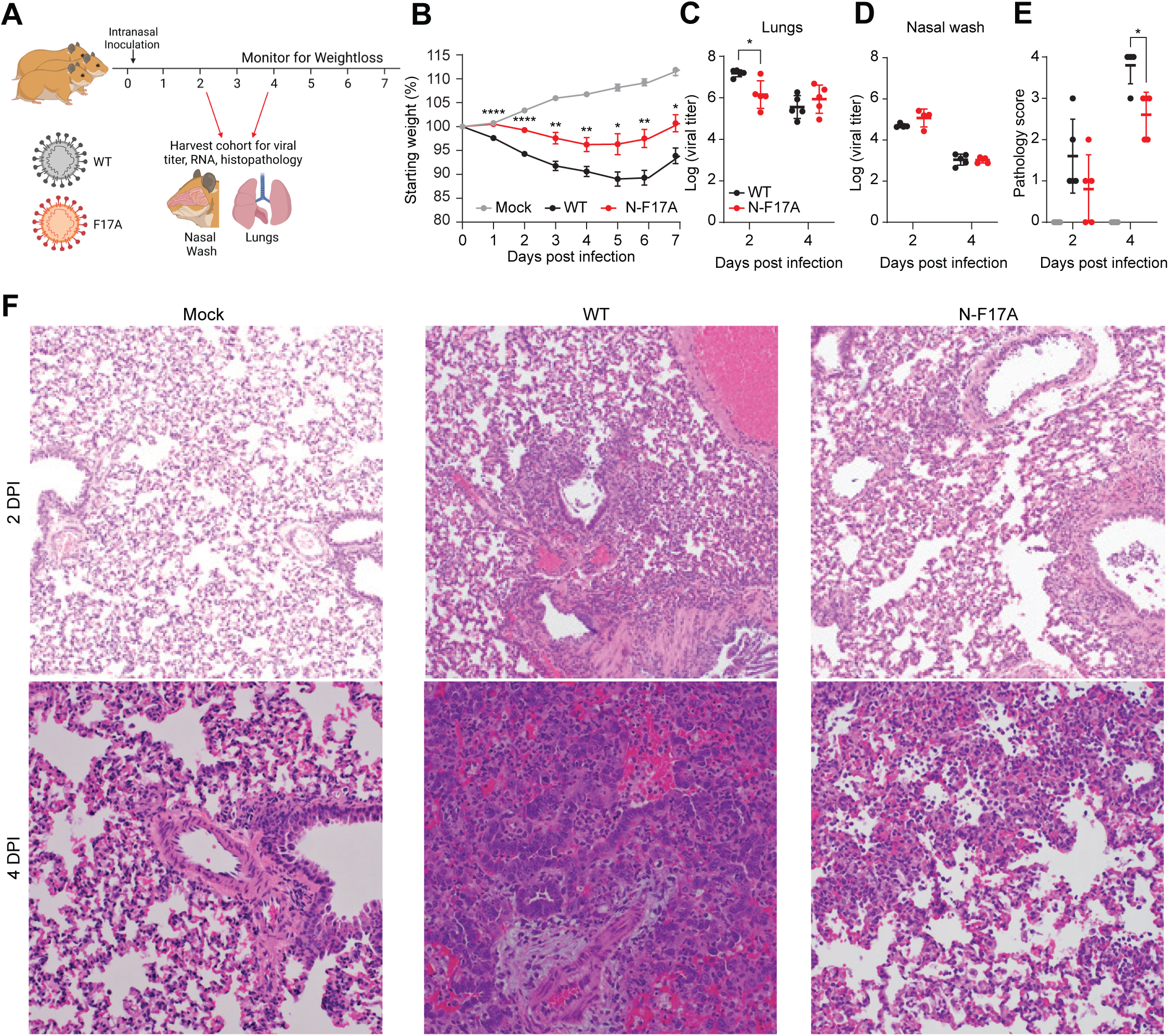
N-F17A Reduces SARS-CoV-2 Replication and Pathogenesis In Vivo. (A) Illustration of hamster infection assay design. Three- to four-week-old male Golden Syrian hamsters (n = 5) were intranasally inoculated with PBS (mock) or PBS containing 10^5^ focus forming units of WT or N-F17A mutant SARS-CoV-2. (B) Weight change of animals for 7 days post infection. Stars indicate the statistical differences betweeen WT and N-F17A mutant. (C-E) At 2 and 4 days post infection, cohorts of 5 animals were nasal washed with PBS and then sacrificed. Viral titers of nasal wash (C) and lung tissue (D) were analyzed by focus forming assay. Lung tissue pathology (E) was analyzed by H&E and scored by a blinded pathologist. (F) Representative H&E staining of lung tissues at 2 and 4 days post infection. Error bars represent mean ± SD in all panels. \**P* < 0.05; \*\**P* < 0.01; \*\*\*\**P* < 0.0001 by two-tailed Student’s t test.

### N protein rewires the G3BP1 network to free viral gRNA from being diverted into condensates

Expression of the SARS-CoV-2 N protein increases as infection progresses, eventually becoming the most abundant viral protein in infected cells ^7–13^. To investigate the effects of different expression levels of N protein, we examined Vero E6-TMPRSS2 cells infected with SARS-CoV-2 and assessed their relative expression of N and G3BP1 inside stress granules. We observed an inverse relationship between N protein expression levels and stress granule formation, wherein cells with low levels of N protein tended to form stress granules, whereas those with high levels of N protein showed a reduced propensity for stress granule formation (**Figure 6A**). We confirmed this inverse correlation by quantitative imaging of 998 infected cells, in which N protein expression levels were measured by antibody staining intensity and stress granules were measured by the enrichment ratio of G3BP1 inside eIF3η-positive puncta (**Figure 6B**).

**Figure 6.**
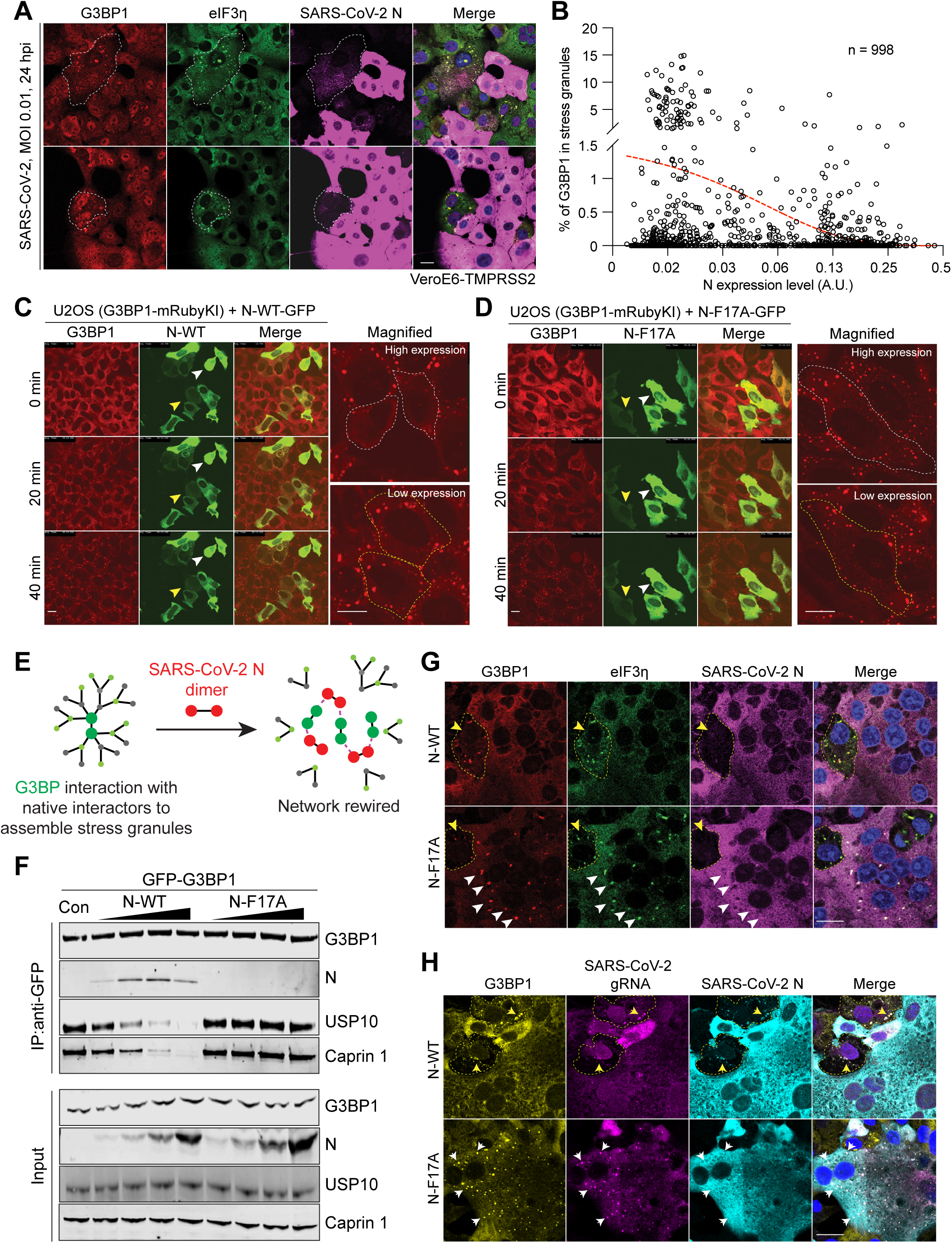
N Protein Rewires the G3BP1 Network to Free Viral gRNA From Being Diverted into Condensates. (A-B) Vero E6-TMPRSS2 cells were infected with SARS-CoV-2 at MOI of 0.01 for 24 h and fixed for staining. Images of 998 infected cells were analyzed quantitatively. N protein expression levels were measured by antibody staining intensity and stress granule formation was measured by the enrichment ratio of G3BP1 inside eIF3η-positive puncta (B). The red dotted trendline indicates the inverse correlation between N protein expression level with G3BP1 enrichment inside stress granules. (C-D) G3BP1-tdTomato-KI U2OS cells were transfected with GFP-tagged N-WT or N-F17A. Stress granule formation was induced by 500 μM sodium arsenite and monitored with live cell imaging for 1 h. Representative images are shown 0, 20 and 40 min post treatment. Yellow and white arrowheads indicate low- and high-expressing cells, respectively, as shown in higher magnification images. (E) Illustration of how N protein may rewire the G3BP1-centered protein interaction network. (F) HEK293T *G3BP1/2* DKO cells were transfected with G3BP1-GFP and lysed after 48 h. Increasing amounts of purified N-WT and N-F17A proteins were added to the cell extracts, which were further captured with magnetic beads conjugated with GFP antibody for IP. Bound proteins were analyzed by immunoblot. (G) Vero E6-TMPRSS2 cells were infected with WT or N-F17A mutant SARS-CoV-2 at MOI of 0.01. Cells were fixed for staining at 24 hpi. (H) Vero E6-TMPRSS2 cells were infected with WT or N-F17A mutant SARS-CoV-2 at MOI of 0.01. At 24 hpi, cells were stained for viral gRNA, G3BP1, and N. Scale bar, 20 μm.

To determine the extent to which expression of the N protein, among many other changes in infected cells, and its interaction with G3BP1, was specifically suppressing stress granule formation, we expressed GFP-tagged N protein in G3BP1-tdTomato-KI U2OS cells and induced stress granule formation using sodium arsenite. Consistent with a direct role of N protein in suppressing stress granule formation, we observed impaired stress granule formation in cells with high expression of N-WT-GFP, but not in cells with high expression of N-F17A-GFP (**Figure 6C, D**). In cells with moderate or low expression of N-GFP, N-WT and N-F17A appeared to be recruited into stress granules, and we observed no significant impairment in stress granule formation (**Figure 6C, D**). These results suggest that high levels of N protein expression suppress stress granule formation, and this effect requires interaction between G3BP1 and the N protein. Thus, we hypothesized that increasing N protein expression during SARS-CoV-2 infection competes with native G3BP1-interacting proteins for binding to G3BP1. This competitive binding by the N protein “rewires” the G3BP1-centered network and thereby inhibits stress granule formation (**Figure 6E**). Indeed, co-IPs from cells demonstrated that G3BP1 increasingly lost interactions with USP10 and caprin 1 with increasing amount of N-WT protein, but not with N-F17A (**Figure 6F**). Consistent with this result, even at a late stage of infection when the N protein was highly expressed and cells began to exhibit syncytium morphology, the formation of stress granules was still observed in cells infected with mutant SARS-CoV-2 (N-F17A), but not WT SARS-CoV-2 (**Figure 6G**).

In stress granules induced by environmental stress, longer mRNAs, which provide more binding valency for G3BP1/2 and other RNA-binding proteins ^20^, are preferentially enriched inside stress granules compared to shorter mRNAs ^80^. Applying this concept to stress granules formed during SARS-CoV-2 infection, we hypothesized that longer viral mRNAs, especially the ∼30kb long viral gRNA, would be diverted into stress granules. To test this hypothesis, we co-stained G3BP1 and viral gRNA, which we labeled using a mixture of probes targeting different conserved regions of ORF1ab ^81^. In cells infected with wild type SARS-CoV-2, we observed that stress granules showed enrichment of viral genomic RNA in about 55% cells (125 of 227 cells). The failure of detecting gRNA signal in stress granules of the other 45% of cells might due to that cells were still in the early stage of infection with low levels of gRNA (**Figure 6H**). Remarkably, in cells infected with mutant SARS-CoV-2 (N-F17A), which showed persistent stress granules even in later stages of infection when gRNA was abundant, we also saw an enrichment of viral gRNAs inside stress granules (**Figure 6H**). The diversion of viral gRNAs, and perhaps other long viral RNAs as well, into stress granules, may have several antiviral consequences. For example, the stress granule may represent an environment in which viral RNA remains inaccessible for translational machinery and therefore decreases viral replication. As stress granules have been proposed to serve as platforms that enhance the innate immune response ^82^, the presence of long viral RNA in these assemblies may also trigger antiviral signaling cascades. Thus, SARS-CoV-2 may use the interaction between G3BP1/2 and N protein to disassemble stress granules and promote viral replication, a function disrupted by SARS-CoV-2 (N-F17A), which lost interaction with G3BP1/2 and replicated less well than WT virus.

## Discussion

In this study, we used biochemical and structural approaches to define the precise basis for the interaction between G3BP1/2 and SARS-CoV-2 N protein. We first demonstrated that these proteins directly interact via the NTF2L domain of G3BP1 and aa 1-25 of the N protein. When we compared this interaction to other G3BP1 interactors that utilize the same NTF2L binding pocket, we found a feature unique to the G3BP1-N interaction that enabled us to selectively impair the G3BP1-N interaction while leaving other host interactions intact. Specifically, we demonstrated that both ancestral and variant forms of N protein bind to G3BP1 by inserting the side chain of N-F17 into the center of a hydrophobic binding pocket of the NTF2L domain. Other known NTF2L interactors, including USP10, caprin 1, Nup_FxFG and SFV-nsP3, also inserted a phenylalanine side chain into this binding pocket, although the secondary interaction surfaces differed from those used by the N protein. Based on these studies, we engineered specific point mutations in the N protein (F17A) and G3BP1 (V11A and F124W) that specifically and reciprocally disrupt their binding. Introducing the N-F17A mutation into the SARS-CoV-2 genome resulted in attenuation of viral replication and pathogenesis both in vitro and in vivo. Together, these data demonstrate the importance of the G3BP1-N interaction in disrupting stress granule formation and driving SARS-CoV-2 infection and pathogenesis.

G3BP1/2 acts as surveillance machinery in the host cell cytoplasm, where increasing concentrations of uncoated mRNAs trigger the condensation of RNAs with proteins into stress granules ^20^. Typically, the source of these uncoated mRNAs is a shutdown in cellular translation induced by either environmental stresses or viral infection, and consequent ribosome runoff from polysomes. When the process of condensation is triggered, G3BP1/2 serves as a hub for a large number of stress granule proteins that together create a network of interactions to promote condensation and the formation of a phase-separated RNP granule. The vast majority of these interactions occur in a mutually exclusive manner via a common versatile binding pocket in the NTF2L domain of G3BP1/2. We found that the N protein exploits the critical nature of this interaction surface by filling the NTF2L binding pocket and thereby impairing the ability of G3BP1 to form the stress granule interaction network. Remarkably, the mode of binding of N protein with G3BP1 includes interaction of N-F17 and flanking amino acids with G3BP1 V11 and F124 within the NTF2L binding pocket. This observation afforded us the opportunity to genetically engineer a version of N that loses interaction with G3BP1 while retaining the interaction between G3BP1 and other host proteins, obviating the need for genetic knockout or knockdown studies that can be confounded by compensatory responses or pleiotropic effects.

How does the assembly and disassembly of stress granules relate to the replication of SARS-CoV-2? We noted the formation of transient stress granules in early stages of infection, a phenomenon also commonly observed in cells infected by other RNA viruses ^82^. These transient stress granules rapidly disassembled as infection proceeded and the expression level of N protein increased. Furthermore, we observed increased formation of persistent stress granules in cells infected with SARS-CoV-2 N-F17A, a form of the virus that is specifically unable to interact with G3BP1. These stress granules were enriched with viral gRNA, indicating that the long (∼30 kb) gRNA of SARS-CoV-2 is diverted into stress granules. Indeed, longer mRNAs, which provide more binding valency for G3BP1/2 and its interacting RNA-binding proteins, trigger condensate formation more easily (i.e., with a lower concentration threshold) and are preferentially enriched inside condensates compared with shorter mRNAs ^80^. Incorporation of long viral RNAs into stress granules may attenuate viral replication by (1) sequestering viral RNA from the cellular machinery required for translation and replication, and/or (2) facilitating the activation of host viral RNA sensors to enhance innate immune responses.

Some evidence suggests that the relationship between SARS-CoV-2 and stress granules may have additional layers of complexity. For example, one recent study demonstrated that overexpression of a peptide derived from SFV-nsP3, which binds the G3BP1 NTF2L domain, *suppressed* SARS-CoV-2 replication in Vero E6 cells ^46^. This SFV-nsP3 peptide bound the hydrophobic pocket within G3BP1 NTF2L, outcompeting both the SARS-CoV-2 N protein and host proteins and thus potently suppressing stress granule formation ^83^. These findings are consistent with our observed relationship between the N protein and G3BP1 with respect to phase separation (**Figure S3**). Specifically, we found that although N protein did not undergo phase separation alone at physiological salt concentrations, the addition of purified G3BP1 induced co-condensation with N-WT, but not N-F17A. Moreover, the addition of purified G3BP1 lowered the concentration threshold for co-condensation of N protein and RNA. Thus, at very high levels of N protein expression, as occur during later stages of infection, it is possible that the G3BP1-N interaction may have the opposite effect as it does earlier in infection (**Figure S4**). In this context, G3BP1 may aid in the co-condensation of N protein with viral RNAs. This increased engagement of N protein with viral RNAs may have several pro-viral effects, including increased viral RNA replication and translation, suppression of the innate immune response, and increased packaging of viral gRNA.

Intriguingly, we found a modest reduction in G3BP1-N interaction due to the N-P13L mutation carried by the Omicron variant ^84^ and its sublineages (e.g., BA.1, BA2, BA4, and BA5 ^85^), which are likely to be the genetic background from which new SARS-CoV-2 variants will emerge ^68^. Lower hospitalization and mortality rates have been observed in patients infected with the Omicron variant ^86–89^, effects attributable to protection afforded by widespread vaccines and previous infections ^90^ as well as reduced virulence of Omicron compared to preceding variants ^91^. In our studies, this N-P13L mutation, just four amino acids away from F17, was located in close proximity to the primary interaction surface with the G3BP1-NTF2L domain and modestly reduced the binding affinity of N protein with G3BP1. We look forward to future studies to determine whether the decreased interaction between N-P13L and G3BP1/2 may contribute to the decreased virulence of the Omicron variant.

## Methods

### Cell culture and transfection

HEK293T and U2OS-tdtmato ^20^ cells were cultured in Dulbecco’s modified Eagle’s medium (HyClone) supplemented with 10% fetal bovine serum (HyClone). Vero E6 and Vero E6-TMPRSS2 (XenoTech; JCRB1819) cells were cultured with DMEM (Gibco; 11965–092) supplemented with 10% fetal bovine serum and 1% antibiotic/antimitotic (Gibco; 5240062). Calu-3 2b4 cells were grown in DMEM with 10% defined FBS, 1% antibiotic/antimitotic, and 1 mg/ml sodium pyruvate. Lipofectamine 2000 (Thermo Fisher; 11668019) was used for transient transfections per the manufacturer’s instructions.

### Constructs

Full-length cDNA of N protein was synthesized according to the SARS-CoV-2 genomic reference sequences but with alternative codons (Integrated DNA Technologies). N-FLAG was inserted into the backbone of pEGFP-C3 plasmid (with GFP removed), AcGFP-N was inserted into the backbone of pAcGFP1-C1, and His-SUMO-N was inserted into the backbone of pETite-HisSUMO (Lucigen) using NEBuilder HiFi DNA Assembly Master Mix kit (NEB; E2621). Using the same methods, human G3BP1 cDNA was inserted into pGEX-2T; G3BP1-EGFP, mCherry-nsP3-25mer (449-472) was inserted into the backbone of pEGFP-C3 plasmid (with GFP removed); EGFP-N-25mer (1-25), EGFP-caprin1-21mer (361-381), EGFP-USP10-25mer (1-25), EGFP-nsP3-25mer (449-473) were inserted into the backbone with pEGFP-C3. Truncations of N-FLAG (48-419, 175-419, 210-419, 1-361, 1-267, 1-209), GST-G3BP1 (GST-NTF2L, GST-deltaNTF2L), and AcGFP-N (1-47, 1-35, 1-25) were created using NEBuilder HiFi DNA Assembly Master Mix kit. Point mutations of N-FLAG (F17A and F17W), AcGFP-N (F17A), GFP-tagged peptides (EGFP-N-25mer-F17A, EGFP-caprin1-21mer-F372A, EGFP-USP10-25mer-F10A and EGFP-nsP3-25mer-F451/468A), GST-NTF2L (F33W), G3BP1-GFP (V11A, F112A, F33A, F33W, F15A, F15W, F124A, F124W) were created by the Q5 Site-Directed Mutagenesis kit (NEB; E0554S).

### Generation of HEK293T *G3BP1/2* DKO cell line

To generate HEK293T *G3BP1/2* DKO cells, gRNAs for G3BP1 and G3BP2 were designed through http://crispr.mit.edu/ and synthesized DNA oligos were ligated into BbsI-(NEB) digested px459 vectors (Addgene; 62988) ^92^. Parental HEK293T cells were transiently transfected with gRNA vector for 2 days, followed by the addition of 2 μg/ml puromycin for 3 days. Knockout clones were obtained by single-cell dilution in 96-well plates and successful knockout was verified by antibody staining.

### Immunoblotting

Cells were washed twice with PBS and lysed with RIPA buffer (25 mM Tris-HCl, pH 7.6, 150 mM NaCl, 1% NP-40, 1% sodium deoxycholate, 0.1% SDS; Thermo Fisher; 89901) supplemented with 1 mM EDTA (Invitrogen; 15575020) and proteinase inhibitor cocktail (Roche; 1183617001). Lysates were centrifuged for 15 min at 4°C at 20,000 x g. 4× NuPAGE LDS sample buffer (Thermo Fisher; NP0007) was added to the supernatant. The samples were boiled at 90°C for 5 min, separated in 4 to 12% NuPAGE Bis-Tris gels (Thermo Fisher; NP0336BOX or NP0321BOX) and transferred to PVDF membranes (Thermo Fisher; IB24001) using an iBlot 2 transfer device (Thermo Fisher). Membranes were blocked with Odyssey blocking buffer (LI-COR Biosciences; 927–50000) and then incubated with primary antibodies at 4°C overnight. Primary antibodies used in this study were anti-G3BP1 (mouse; BD Biosciences; 611127), anti-G3BP1 (rabbit; Proteintech; 13057-2-AP), anti-G3BP2 (rabbit; Thermo Fisher; PA5-53776), anti-N (mouse; Sino Biological; 40143-MM05), anti-N (rabbit; Sino Biological; 40143-R019), anti-USP10 (Rabbit; Proteintech; 19374-1-AP), anti-caprin 1 (Rabbit; Proteintech; 15112-1-AP), anti-GST (rabbit; BioVision; 3997-30T), anti-FLAG M2 antibody (mouse; Sigma-Aldrich; F1804), anti-mCherry (Rabbit; BioVision; 5993), anti-Hisx6 tag (mouse; abcam; ab18184), anti-GFP (Rabbit, Cell Signaling Technology; 2555). Membranes were washed three times with TBS-T (0.05% Tween) and further incubated with IRDye 680RD/800CW-labeled secondary antibodies (LI-COR Biosciences; 926–68073 or 926–32212) at a dilution of 1:10,000. Membranes were visualized with an Odyssey Fc imaging system (LI-COR Biosciences).

### Immunoprecipitation

Cells were washed twice with PBS and lysed with IP lysis buffer (25 mM Tris-HCl, pH 7.4, 150 mM NaCl, 1% NP-40, 10% glycerol) supplemented with proteinase inhibitor cocktail (Roche; 1183617001). Lysates were centrifuged at 4°C for 15 min at 20,000 x g. Supernatants were incubated with anti-FLAG M2 magnetic Beads (Millipore; M8823) or GFP-Trap magnetic beads (ChromoTek; gtma-10) at 4°C overnight. Beads were washed three times with lysis buffer, with the second wash supplemented with 0.1 mg/ml RNaseA (Thermo Fisher; EN0531) to remove RNA and hence the RNA-mediated protein-protein interactions. The beads were boiled in 1× NuPAGE LDS sample buffer at 70°C for 5 min and analyzed by immunoblotting.

### Protein interactome by LC-MS/MS

Proteins interacting with GFP-N-WT, GFP-N-F17A, G3BP1-GFP WT, and G3BP1-GFP mutants were immunoprecipitated as described above. Resulting samples were isolated from gels and digested with trypsin overnight. Samples were loaded on a nanoscale capillary reverse-phase C18 column by an HPLC system (Thermo Ultimate 3000) and eluted by a gradient (∼90 min). Eluted peptides were ionized by electrospray ionization and detected by an inline mass spectrometer (Thermo Orbitrap Fusion). Database searches were performed using SEQUEST search engine using an in-house SPIDERS software package. Tandem mass spectrometry spectra were filtered by mass accuracy and matching scores to reduce the protein false discovery rate to ∼ 1%. The total number of spectral counts for each protein identified was reported by sample. The p value of spectral count changes between different groups was derived by G-test ^93^. Proteins were accounted as confident N protein interactors if they were repeatedly found in all four independent assays. A composite protein-protein interaction (PPI) network was built by combining STRING ^94^. For the comparison of two interactomes, proteins with spectral count >2, at least twofold change, and the p-value less than 0.05 were considered as significant changes.

### RNA cross-linking immunoprecipitation

The binding of N-WT and N-F17A with cellular RNA was tested as previously described ^20^. Briefly, HEK293T cells transfected with pAcGFP-N-WT or pAcGFP N-F17A were washed with PBS and exposed to UV (254 nm, 400 mJ/cm^2^ followed by 200 mJ/cm^2^). Cells were then harvested in lysis buffer containing 20 mM Tris HCl, pH 7.5, 137 mM NaCl, 1% Triton X-100, 2 mM EDTA, 1x protease inhibitor and incubated for 10 min on ice. Lysates were then successively treated with 8 U/mL DNase I (NEB; M0303S) for 5 min at 37°C and 4 U/mL RNase I for 3 min at 37°C. The reaction was stopped by the addition of 0.4 U/μL RNaseIn (Promega; N2615). Lysates were then centrifuged at 21,000 x g for 10 min at 4°C. The supernatant fraction was incubated with 10 μl GFP-Trap magnetic beads at 4°C overnight. The beads were washed twice with lysis buffer, twice with 1 M NaCl, and twice with lysis buffer again. Beads were further suspended in 100 μl 10 mM Tris-HCl, pH 7.5, and treated with 2 units of calf intestinal alkaline phosphatase (Promega; M182A) at 37°C for 10 min at 1000 rpm in a Thermomixer. Beads were washed with lysis buffer twice and RNA labeling was performed with an RNA 3′ End Biotinylation Kit (Pierce; 20160). After washing with lysis buffer twice, beads were boiled in 1x LDS sample buffer (Life Technologies; NP007) with 100 mM DTT. Protein-RNA complexes were separated by 4–12% NuPAGE Bis-Tris gels, transferred to nitrocellulose membranes, and blotted with IRDye 680LT streptavidin (LI-COR; 926–68031).

### Protein purification

GST-G3BP1 full-length and mutants, His-SUMO-N-WT and His-SUMO-N-F17A were expressed and purified from *E. coli* Rosetta 2(DE3) cells (Millipore; 71400-4). *E. coli* transformed with indicated expression plasmids were grown to OD_600_ of 0.8 and induced with 1 mM IPTG at 16°C overnight. Pelleted cells were resuspended in lysis buffer (300 mM NaCl, 50 mM HEPES 7.5, 1 mM DTT, protease inhibitor). After sonication, lysates were pelleted at 30,000 x g at 4°C for 30 min. For GST-tagged G3BP1 proteins, supernatants were applied to packed Glutathione-Sepharose 4B resin (Cytiva; 17-0756-01) and were eluted with 10 mM glutathione (Sigma-Aldrich; G4251) in lysis buffer. For His-SUMO-tagged proteins, the lysis buffer was supplemented with 60 mM imidazole. Cell lysate supernatants were applied to packed Ni-NTA beads (QIAGEN; 30210). After washing with 20 column volume of lysis buffer, the proteins were eluted with 300 mM imidazole in lysis buffer. All eluted proteins were further treated with 0.1 mg/ml RNaseA to remove RNA ^95^. The fractions were analyzed by SDS-PAGE, pooled, and concentrated. For proteins that needed tags removed, TEV protease ^96^ or SUMO Express proteinase (Lucigen, 30801-2) was respectively incubated with GST-TEV-tagged G3BP1 proteins and His-SUMO-tagged N proteins at 4°C overnight. The proteins were then purified by Superdex 200 16/200 column (GE) equilibrated in SEC buffer (300 mM NaCl, 50 mM HEPES 7.5, 1 mM DTT). The fractions were analyzed by SDS-PAGE, pooled, concentrated, filtered, flash frozen in liquid nitrogen, and stored at −80°C.

### Peptide synthesis

All peptides were synthesized by the Hartwell Center for Bioinformatics and Biotechnology at St. Jude Children’s Research Hospital, Molecular Synthesis Resource, using standard solid-phase peptide synthesis chemistry. The peptides were reconstituted from lyophilized form into DMSO for subsequent experiments.

### Pulldown

GST-tagged proteins (100 nM), including GST-G3BP1-full length, GST-NTF2L, GST-NTF2L-F33W, and GST-ΔNTF2L, were incubated with His-SUMO tagged proteins (100 nM), including His-SUMO-N-WT or His-SUMO-N-F17A, for 1 h at room temperature in binding buffer (50 mM HEPES, pH 7.5, 150 mM NaCl, 1% Triton X-100, 1 mM DTT). Complexes were pulled down using Glutathione-Sepharose 4B resin (Cytiva; 17-0756-01) or Ni-NTA resin (QIAGEN; 30210) for 2 h at 4°C, washed 3 times with binding buffer, with the second wash supplemented with 0.1 mg/ml RNaseA to remove RNA and hence RNA-mediated protein-protein interactions. Beads were boiled in 1× NuPAGE LDS sample buffer at 70°C for 5 min and analyzed by Coomassie blue staining or immunoblotting with anti-GST and anti-6xHis antibodies.

### Crystallography studies

Co-crystals of the NTF2L domain of G3BP1 bound to peptides were obtained by sitting drop at 4°C. Crystals were cryoprotected with 15% glycerol and flash cooled. X-ray diffraction data was collected at the European Synchrotron Radiation Facility (ESRF). Data processing and scaling were performed with DIALS ^97^. The diffraction data was phased by molecular replacement using PDB 5FW5 ^61^ as the search model. The initial model was built by Phaser MR ^98^ and completed with Coot ^99^. Refmac ^100^ was used to improve density between rounds of manual building.

### Surface plasmon resonance

Binding studies were performed at 25 °C using a Biacore T200 optical biosensor (Cytiva). Anti-GST antibodies (Cytiva) were covalently attached to a carboxymethyl dextran-coated gold surface (CM-4 Chip; Cytiva). GST-tagged NTF2L constructs were captured on the flow cell surfaces, and GST was captured on the reference surface to account for any non-specific binding to the GST tag. The peptide analytes were prepared in 20 mM HEPES (pH 7.5), 150 mM NaCl, 5% glycerol, 0.01% Triton X-100, 5% DMSO as three-fold dilution series and injected in triplicate at flow rate 75 µL/min. Data were processed, double-referenced, solvent corrected, and analyzed using the software package Scrubber 2 (Version 2.0c, BioLogic Software). Equilibrium dissociation constants (K_D_) were determined by fitting the data to a 1:1 (Langmuir) binding model.

### Liquid-liquid phase separation

In vitro LLPS experiments were performed at room temperature. Indicated concentrations of protein(s), RNA, or Ficoll 400 were mixed together in low binding tubes (COSTAR; 3206) and transferred to a sandwiched chamber created by cover glass and a glass slide with a double-sided spacer (Sigma-Aldrich; GBL654002). Samples were observed under a DIC microscope using a Leica DMi8 microscope with a 20x objective. All imaged were captured within 5 min after LLPS induction. RNA used in LLPS assays was isolated from HEK293T cells using TRIzol (Thermo Fisher; 15596026) and the concentration of RNA was measured by Nanodrop (Thermo Fisher).

### SARS-CoV-2 N protein amino acid variation analysis

SARS-CoV2 genome sequences (n=8,297,154) deposited in GISIAD ^71^ were accessed in March 2022. N protein residue usage analyses were based on the multiple sequence alignments. Residues used in the N protein of each genome were enumerated and fraction of residue usage was derived via customized scripts.

### SARS-CoV-2 GFP/ΔN trans-complementation cell culture system

The reproduction of transcription and replication-competent virus-like-particles in Caco2-N (WT) and Caco2-N (F17A) cells was tested as previously described ^73^. Briefly, five DNA fragments of SARS-CoV-2 GFP/ΔN, SARS-CoV-2 genome (Wuhan-Hu-1) with the ORF of N gene replaced by GFP cDNA, were amplified by PCR in vitro. During the PCR reaction, the T7 promoter was introduced to the 5′ end of the first fragment, and the designed restriction endonuclease sites (BsaI or BsmBI) were introduced to the corresponding ends of PCR products, which were then used to ligate the neighboring DNA fragments in the right order. The in vitro ligated full-length SARS-CoV-2 GFP/ΔN DNA was used as the template to transcribe the viral RNA with mMESSAGE mMACHINE T7 Transcription Kit (Thermo Fisher; AM1344). The mixture of 20 μg of viral RNA and 20 μg N mRNA was added to a 4-mm cuvette containing 0.4 mL of Caco-2-N cells (8×10^6^) in Opti-MEM. A single electrical pulse was given with a GenePulser apparatus (Bio-Rad) with setting of 270 V at 950 μF. Three days post electroporation, P0 virus-like-particles were collected. The expression of N-WT or N-F17A in Caco2 cells was accomplished by lentiviral transduction with pLVX plasmid as the transfer vector. Both Caco2-N (WT) and N (F17A) cells were infected with trVLPs at an MOI of 0.05 for 36 hours. The newly reproduced trVLPs in the cell culture supernatant were collected and used to infect fresh Caco-2-N cells. The percentage of GFP-positive cells at 36 hpi was determined by FACS.

### SARS-CoV-2 mutant generation

The WT SARS-CoV-2 sequence was based on the USA-WA1/2020 strain. It was distributed by the World Reference Center for Emerging Viruses and Arboviruses (WRCEVA) and originally isolated by the USA Centers for Disease Control and Prevention ^76^. Recombinant mutant and WT SARS-CoV-2 were created using the cDNA clone described previously ^74,75^. The generation of recombinant SARS-CoV-2 viruses was approved by the University of Texas Medical Branch Biosafety Committee.

### SARS-CoV-2 in vitro infection

Infection of Vero E6 and Calu-3 2b4 cells were performed according to previously published standard protocols ^75,101^. Briefly, cells were infected at a multiplicity of infection (MOI) of 0.01 by removing the growth media and adding virus diluted in PBS. Cells were subsequently incubated for 45 min at 37°C and 5% CO_2_. Cells were then washed three times with PBS and fresh growth media returned. Virus containing growth media was then sampled at the indicated time points, replacing the removed sample with an equal amount of fresh growth media to maintain a consistent total volume. All infections occurred at Biosafety Level 3 (BSL3) facilities at the University of Texas Medical Branch or St. Jude Children’s Research Hospital.

### Focus forming assay

For viral titrations, focus forming assays were performed as described previously ^102,103^. Briefly, samples (cell supernatants, nasal washes, or homogenized lung tissue) underwent 10-fold serial dilutions and were then used to infect Vero E6 cells seeded to 96-well plates. After a 45-min incubation at 37°C and 5% CO_2_, 100 μl 0.85% methylcellulose overlay was added. After an additional 24-h incubation, methylcellulose was removed by washing three times with PBS and cells were fixed for 30 min in 10% formalin to inactivate SARS-CoV-2. Cells were then removed from the BSL3 and permeabilized with 0.1% saponin/0.1% bovine serum albumin in PBS. Cells were then stained with an anti-SARS-CoV-1/2 nucleocapsid primary antibody (Cell Signaling Technology; 68344) followed an Alexa Flour 555 fluorescent anti-mouse secondary antibody (Invitrogen; A28180). Each well was then imaged with a Cytation 7 cell imaging multi-mode reader (BioTek) and foci counted manually.

### Hamster infections

For hamster experiments, three- to four-week-old male Golden Syrian hamsters were purchased from Envigo. Animals were intranasally inoculated with 100 μL PBS (mock) or PBS containing 105 focus forming units (ffu) of WT or F17A mutant SARS-CoV-2. Animals were monitored for weight loss and signs of disease for 7 days. On 2 and 4 days post infection (dpi), cohorts of 5 animals were nasal washed with PBS and then sacrificed. Lung tissue was then taken to analyze viral titer by focus forming assay and pathology by hemoxylin and eosin (H&E) staining. H&E staining was performed by the University of Texas Medical Branch Histology Laboratory and then analyzed and scored by a blinded pathologist.

All animal studies were carried out in accordance with a protocol approved by the University of Texas Medical Branch Institutional Animal Care and Use Committee and complied with United States Department of Agriculture guidelines in a laboratory accredited by the Association for Assessment and Accreditation of Laboratory Animal Care. Procedures involving infectious SARS-CoV-2 were performed in the Galveston National Laboratory animal biosafety level 3 (ABSL3) facility.

### Immunofluorescence and microscopy

Cells were seeded on the coverslips pre-coated with collagen (Advanced Biomatrix, 5005) in a 24-well plate. For antibody staining, fixed cells were permeabilized with 0.2% Triton X-100 in PBS for 10 min, and blocked with 1% BSA for 1 h. Samples were further incubated with primary antibodies, including antibodies anti-eIF3η (goat, Santa Cruz, sc-16377), anti-G3BP1 (rabbit; Proteintech; 13057-2-AP) and anti-N (mouse; Sino Biological; 40143-MM05), overnight at 4°C, then washed 3 times with PBS and incubated with host-specific Alexa Fluor 488/555/647 (Thermo Fisher) secondary antibodies for 1 h at room temperature. Nuclei was stained with DAPI in PBS (1:10,000, Biotium, 40009). Cell boundaries were stained with Phalloidin cf532 in PBS (1:500, Biotium, 00051). Samples were mounted in ProLong Glass Antifade mounting medium (Thermo Fisher; P36980). Images were captured using a Leica TCS SP8 STED 3X confocal microscope with a 63x oil objective.

For co-visualizing the SARS-CoV-2 genomic RNA together with proteins, FISH staining was performed before the antibody staining. The FISH protocol was adopted from Rensen et al.^81^ with modifications. In brief, a mixture of SARS-CoV-2 ORF1ab targeting primary probes with the FLAP sequences attached were prehybridized with Cy5-labeled secondary probes with a complementary sequence to FLAP. The hybridization reaction contained 40 pmol of primary probes and 50 pmol of secondary probes in 1X NEBuffer buffer (NEB; B7003S), and was carried out in a PCR machine with the cycles of 85°C for 3 min, 65°C for 3 min, and 25°C for 5 min. Fixed cells were washed twice with wash buffer A (2X SSC, Ambion; AM9770) for 5 min, followed by two 5-min washing steps with washing buffer B (2X SSC and 10% formamide, Ambion; AM9342). In a humidified chamber, cells were placed on parafilm and incubated at 37°C overnight with 2 μl prehybridized FISH probes diluted in 100 μl of hybridization buffer, which contained 10% dextran (Sigma-Aldrich; D8906), 10% formamide, and 2X SSC. Cells were washed in the dark at 37°C for 1 h twice with prewarmed washing buffer B. After two PBS washes, stained cells were fixed with 4% paraformaldehyde in PBS and proceeded to downstream antibody staining as described above.

### Stress granule assembly using time-lapse live cell microscopy

A Yokogawa CSU W1 spinning disk with Nikon Elements software was used in time-lapse live cell imaging. Imaging was taken using a 60× Plan Apo 1.4NA oil objective and Perfect Focus 2.0 (Nikon) was engaged for the duration of the capture. During imaging, cells were maintained at 37°C and supplied with 5% CO_2_ using a Bold Line Cage Incubator (Okolabs) and an objective heater (Bioptechs). Stress granule formation in cells was monitored for 60 mins after the addition of 500 μM sodium arsenite.

### Enrichment of G3BP1 in stress granules

The enrichment percentage of G3BP1 protein in stress granules was quantified by CellProfiler as previously described ^104^ with minor modifications. Briefly, the boundaries of the eIF3η-positive puncta and the whole cell were identified using CellProfiler. The ratios of G3BP1 antibody staining signal intensity in the eIF3η-positive puncta (stress granules) to that in the entire cell were used as indicators for the enrichment percentages of G3BP1.

### Statistics and graphs

One-way ANOVAs and two-tailed Student’s t test were calculated in GraphPad Prism. Data distribution was assumed to be normal, but this was not formally tested. Graphs for the analysis were made in Microsoft Excel and GraphPad Prism. All errors corresponding to the standard deviation or standard error of the population are described in figure legends. Statistically significant differences and the number of samples analyzed for each experiment are indicated in figure legends.

## Author Contributions

J.P.T. conceived and supervised the project. Z.Y., B.A.J., V.A.M., X.J., P.Z., M.P.H., J.W., K.P.K., T.C.C., G.W., J.H., J.D., K.W., R.L., K.G.L., R.E.A., P.A.C., D.H.W., K.S.P., J.A.P., S.C.W., M.B.W., H.J.K., R.M., S.S.C., Q.D., V.D.M., and J.P.T. designed and/or performed the experiments and analyzed data. Z.Y., B.A.J., H.J.K., V.D.M., and J.P.T. wrote the primary draft of the manuscript, and all authors contributed to the final version.

## Acknowledgments

We thank Natalia Nedelsky for editorial assistance, Michael Brett Waddell for assistance with SPR, the CAGE at St. Jude Children’s Research Hospital for assistance with Cas9/CRISPR-modified cell lines, and the Center for Applied Bioinformatics for assistance with bioinformatic analysis. This work was supported by grants from HHMI and NIH (R35NS097974) to J.P.T. and from St. Jude Children’s Research Hospital.

---

**Figure S1.**
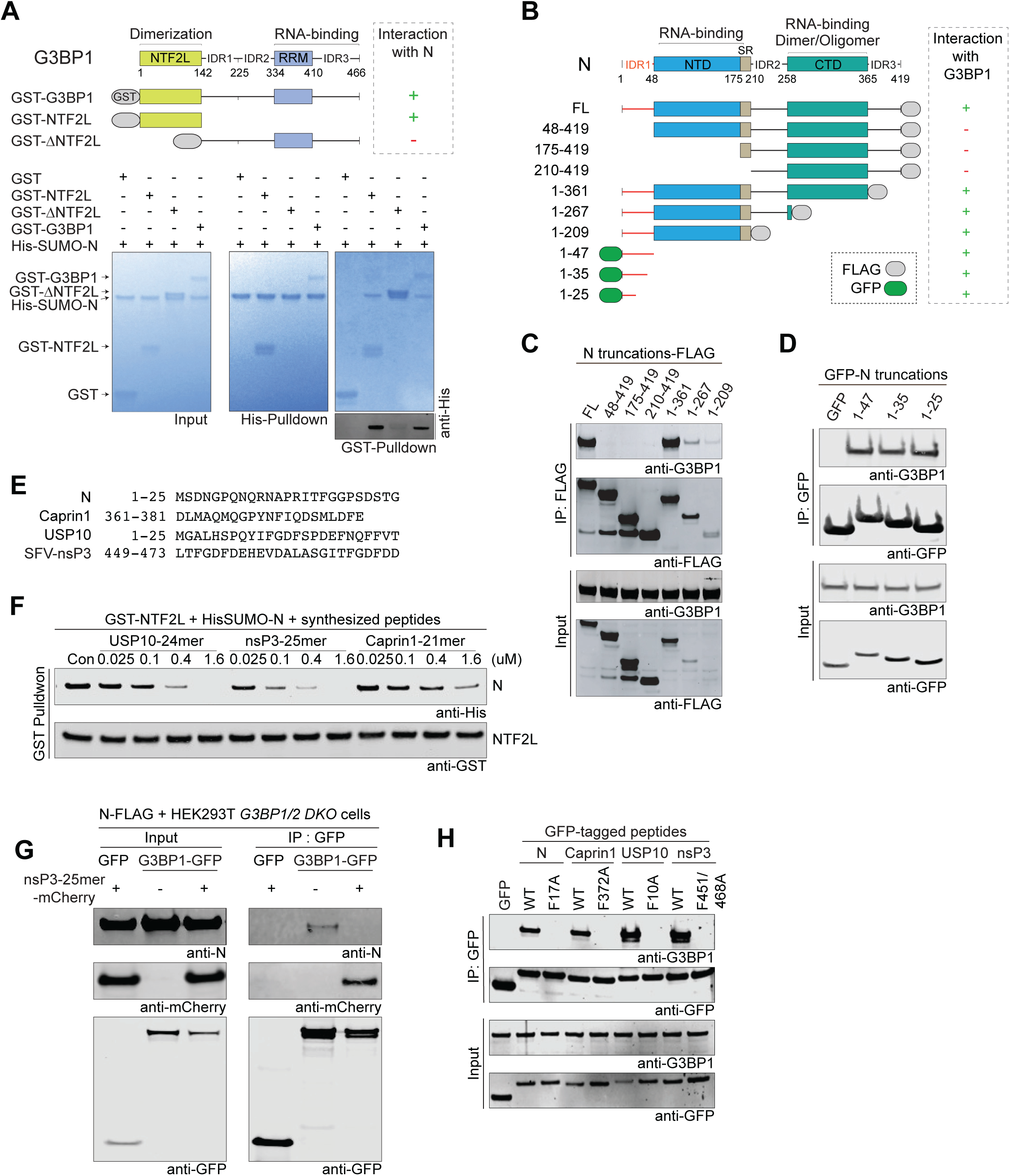
N_1-25_ Directly Interacts with G3BP1-NTF2L Domain and Competes with Other NTF2L Interactors. (A) Top, domain organization of G3PB1 proteins. Bottom, in vitro pulldown assays with purified His-SUMO-N protein and indicated GST-G3BP1 proteins (full-length, NTF2L, ΔNTF2L). Proteins were visualized by Coomassie Blue staining (blue gels) and anti-His immunoblot (grayscale image). (B) N protein truncations used to assess G3BP1 binding in (C) and (D). (C) HEK293T cells were transfected with indicated FLAG-tagged N protein truncations. Cell extracts were captured with magnetic beads conjugated with FLAG antibody for IP and bound proteins were analyzed by immunoblot. (D) HEK293T cells were transfected with GFP-tagged N protein fragments. Cell extracts were captured with magnetic beads conjugated with GFP antibody for IP and bound proteins were analyzed by immunoblot. (E) Amino acid sequences of synthesized G3BP interacting peptides derived from N, caprin 1, USP10, and SFV-nsP3. (F) GST pulldown assay with purified GST-NTF2L (100 nM) and His-SUMO-N (100 nM) with increasing concentrations of caprin 1, USP10, or nsP3 peptides. (G) HEK293T *G3BP1/2* DKO cells were transfected with indicated constructs. Cell extracts were captured with magnetic beads conjugated with GFP antibody for IP and bound proteins were analyzed by immunoblot. (H) HEK293T cells were transfected with GFP-tagged NTF2L interacting peptides. Cell extracts were captured with magnetic beads conjugated with GFP antibody for IP and bound proteins were analyzed by immunoblot.

**Figure S2.**
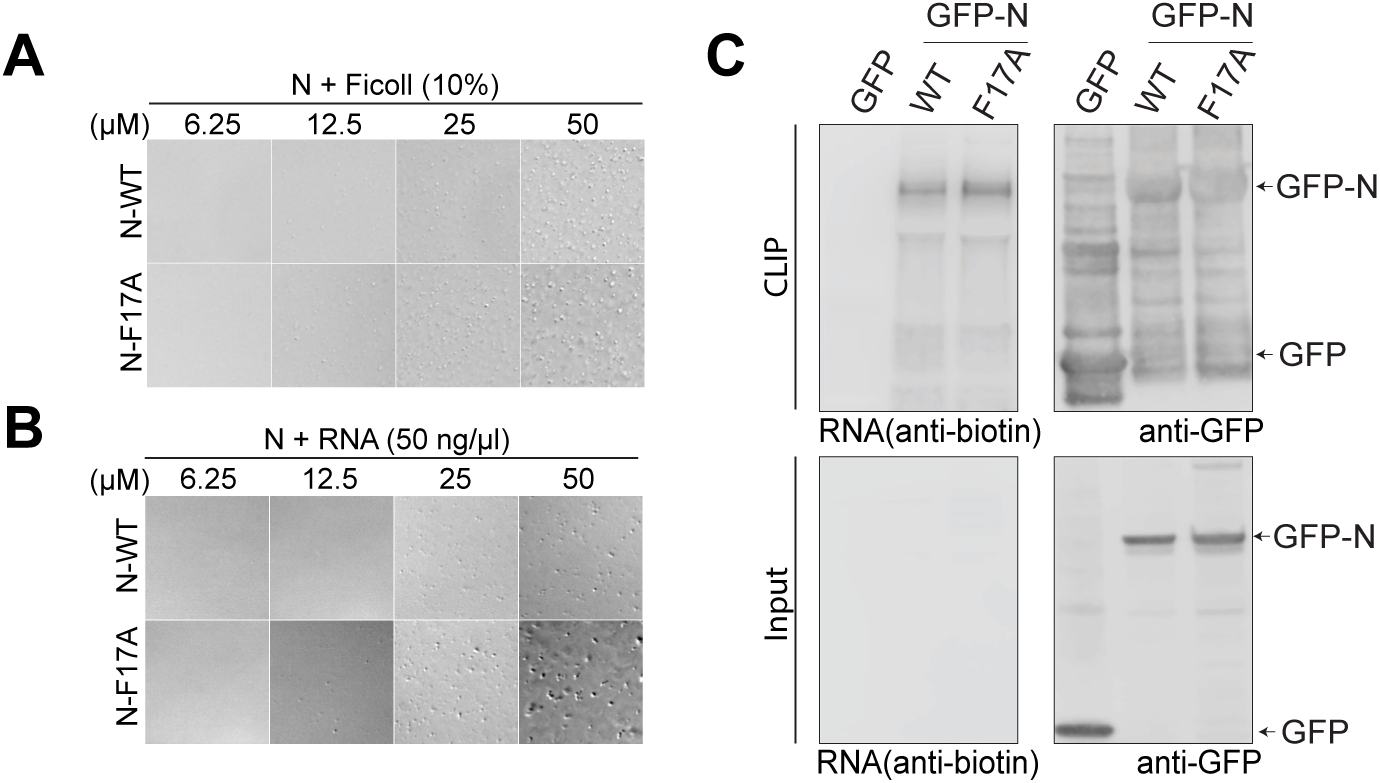
N-F17A Does Not Alter the In Vitro Phase Separation Behavior and RNA Binding of N protein. (A-B) LLPS of purified N-WT and N-F17A with Ficoll (10%) (A) or 50 ng/μl RNA (B) from HEK293T cells. (C) CLIP analysis of GFP-N-WT and GFP-N-F17A transiently expressed in HEK293T cells. Immunoprecipitated N protein cross-linked to RNA was assessed by immunoblotting for biotin (RNA) and GFP (N protein)

**Figure S3.**
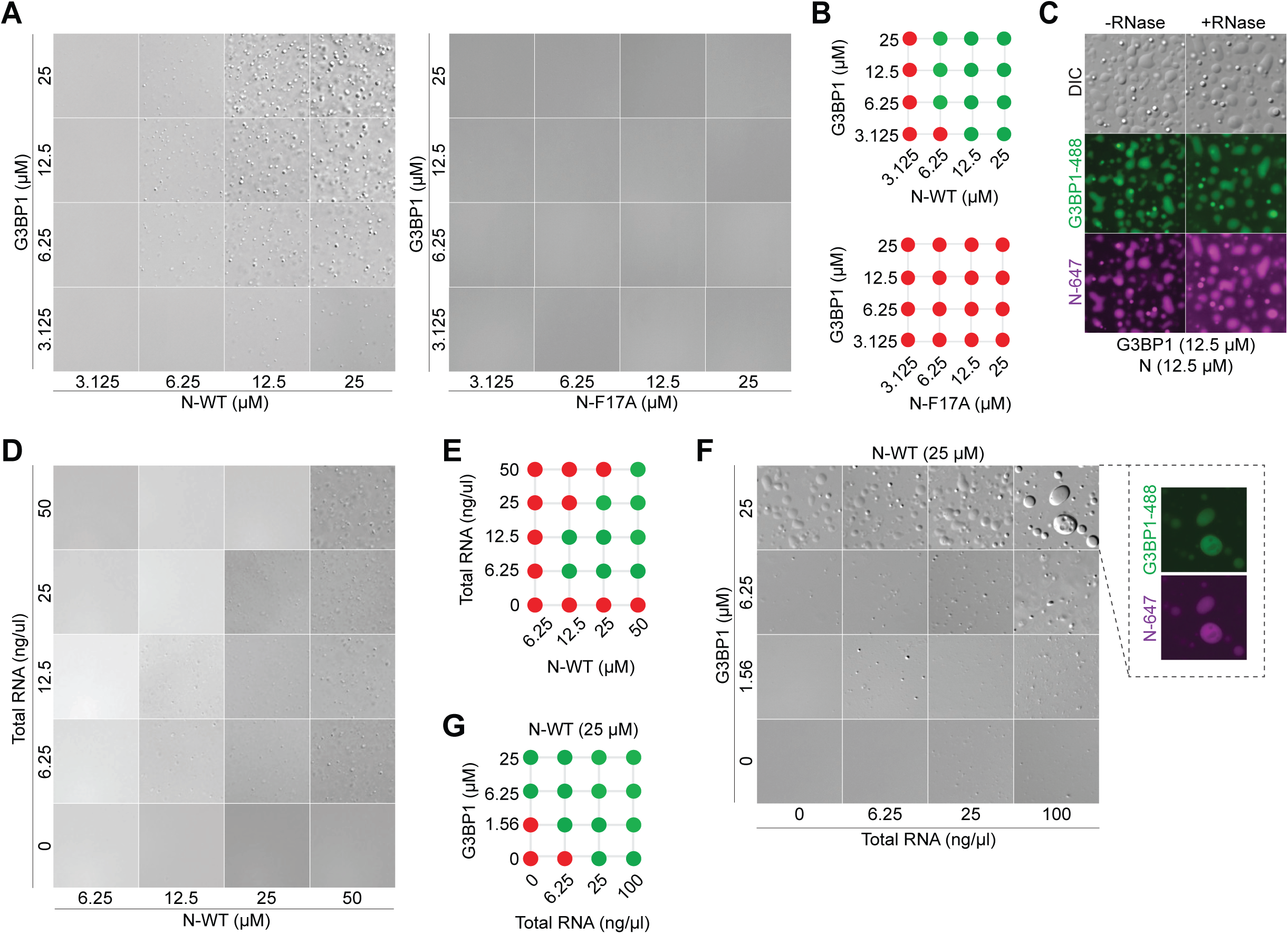
G3BP1 Facilitates N Protein Condensation with RNA. (A, B) LLPS of purified G3BP1 with N-WT or N-F17A proteins, in 150mM NaCl. (C) LLPS of purified G3BP1 (labeled with Alexa Fluor 488) with N (labeled with Alexa Fluor 647), in 150mM NaCl, with or without RNase A (10ug/ml). (D, E) LLPS of purified N with RNA from HEK293T cells. (F, G) LLPS of purified N-WT (25uM) with G3BP1 and RNA from HEK293T cells.

**Figure S4.**
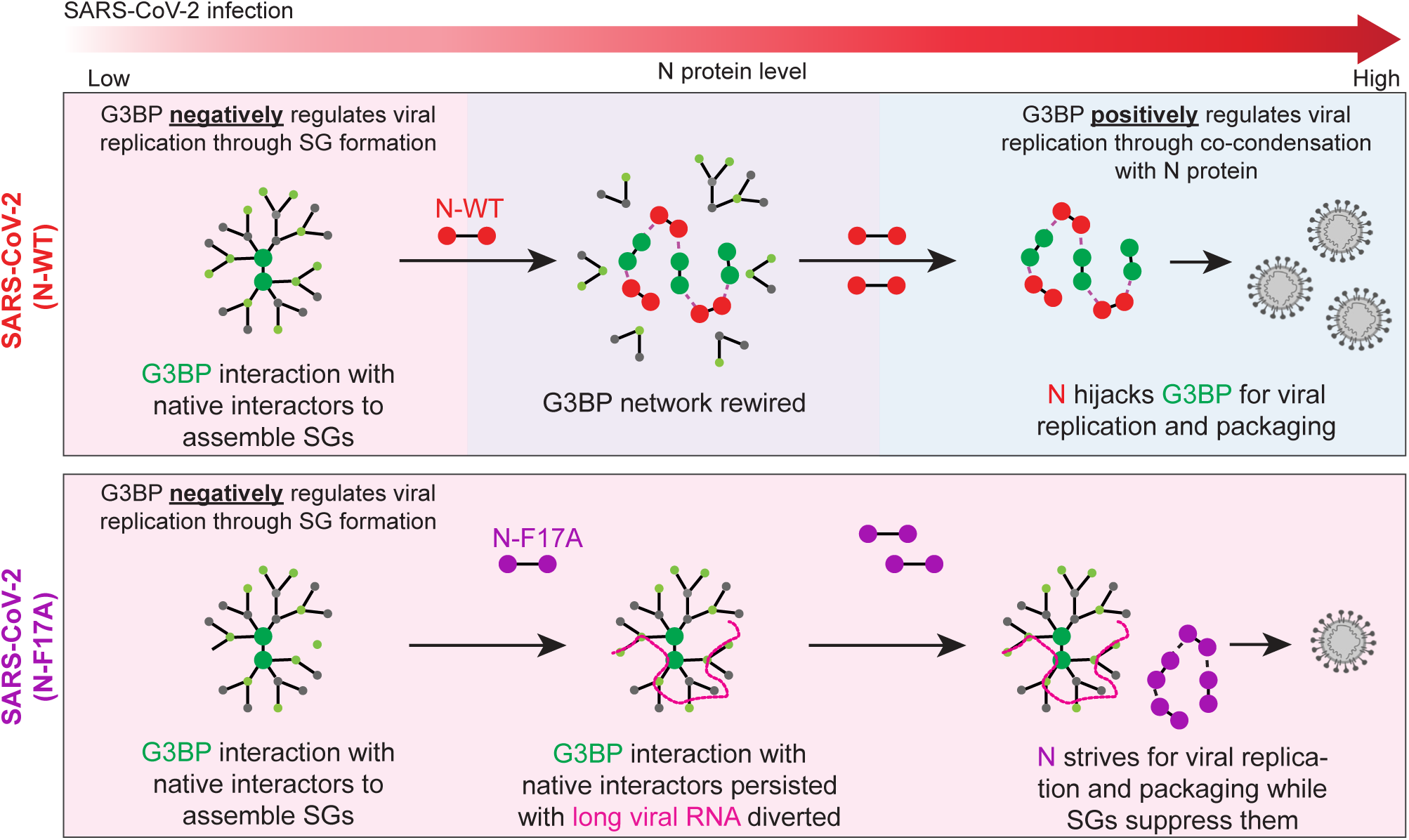

## Notes

### Competing Interest Statement

JPT is a consultant for Nido Biosciences.

